# Tuning of cortical color mechanism revealed using steady-state visually evoked potentials

**DOI:** 10.1101/2023.12.10.570997

**Authors:** Dylan J. Watts, Ana Rozman, Lucy P. Somers, Bora Gunel, Chris Racey, Katie Barnes, Jenny M. Bosten

**Author notes:** Joint first authors (Dylan J. Watts, Ana Rozman).

## Abstract

Color information is thought to be received by the primary visual cortex via two dominant retinogeniculate pathways, one signals color variation between teal and red, and the other signals color variation between violet and lime. This representation is thought to be transformed in the cortex so that there are a number of different cell populations representing a greater variety of hues. However, the properties of cortical color mechanisms are not well understood. In four experiments, we characterized the tuning functions of cortical color mechanisms by measuring the intermodulation of steady-state visually evoked potentials (SSVEPs). Stimuli were isoluminant chromatic checkerboards where odd and even checks flickered at different frequencies. As hue dissimilarity between the odd and even checks increased, the amplitude of an intermodulation component (I_1_) at the sum of the two stimulus frequencies decreased, revealing cortical color tuning functions. In Experiment 1 we found similar broad tuning functions for ‘cardinal’ and intermediate color axes, implying that the cortex has intermediately tuned color mechanisms. In Experiment 2 we found similar broad tuning functions for ‘checkerboards’ with no perceptible edges because the checks were formed from single pixels (∼0.096°), implying that the underlying neural populations do not rely on spatial chromatic edges. In Experiment 3 we manipulated check size and found that color tuning functions were consistent across check sizes used. In Experiment 4 we measured full 360° tuning functions for a ‘cardinal’ cortical color mechanism and found evidence for opponent color responses. The observed cortical color tuning functions were consistent with those measured using psychophysics and electrophysiology, implying that tracking intermodulation using SSVEPs provides a useful method for measuring them.

## 1.0 Introduction

Decades of research in the neural mechanisms of human color vision have produced a relatively good understanding of the representation of color at a receptor level, and in the retinogeniculate color pathways. However, the cortical mechanisms underlying color perception are much less well understood, and though color representation in early cortical areas has been studied using electrophysiology in animal models, there has been relatively little research on the properties of cortical color mechanisms in humans measured directly using neuroimaging. In particular, there is a debate over how far the separation of color information into distinct retinogeniculate pathways remains segregated in the cortex (Schluppeck and Engel, 2002; Gegenfurtner, 2003; Kaneko et al., 2020; Nunez et al., 2021), and over the number and tuning of cortical color mechanisms (Gegenfurtner, 2003; Emery et al., 2017). We used a novel approach using steady-state visually evoked potentials (SSVEPs) to measure and characterize the tuning functions of cortical color mechanisms in humans.

Normal human color vision begins with the relative activations of three types of retinal cone photoreceptors, sensitive to short (S), medium (M) or long (L) wavelengths of light. Cone signals are then thought to be combined postreceptorally by distinct populations of bipolar cells, setting up ‘cardinal’ color pathways which remain segregated in the retina and LGN and at least as far is the input layers of V1 (Gouras, 1974; Chatterjee and Callaway, 2003; Li et al., 2022). The S/(L+M) pathway carried by small bistratified ganglion cells and though koniocellular layers of the LGN signals color differences between lime green and violet. The L/(L+M) pathway carried by the midget retinal ganglion cells and through parvocellular layers of the LGN signals color differences between teal and red (Solomon and Lennie, 2007; Conway et al., 2018).

To what extent retinogeniculate opponent color representations in early visual cortical areas are retained and to what extent they are transformed is not fully understood. Although early psychophysical evidence was consistent with a ‘cardinal’ representation of color inherited from the retinogeniculate mechanisms (Boynton and Kambe, 1980; Krauskopf et al., 1982; Krauskopf and Farell, 1990), later evidence favored a ‘higher order’ color representation based on color mechanisms responsive to hues that are intermediate between the poles of the retinogeniculate color mechanisms (Krauskopf et al., 1986; Webster and Mollon, 1991; Gegenfurtner and Kiper, 1992; Krauskopf and Gegenfurtner, 1992; Hansen and Gegenfurtner, 2013). There has also been fMRI evidence in support of both a ‘cardinal’ color representation in V1 (Engel et al., 1997) and in support of the existence of intermediately tuned color mechanisms (Goddard et al., 2010; Kuriki et al., 2015).

Studies aiming to measure the tuning functions of color selective cortical neurons have been conducted using indirect psychophysical methods in humans and directly using electrophysiology in primates. Results of psychophysical studies have typically shown broad tuning (Sankeralli and Mullen, 1997; Lindsey and Brown, 2004) especially when ‘off-axis looking’ (stimulus detection by neighboring color mechanisms than the one being measured) is minimized (D’Zmura and Knoblauch, 1998; Hansen and Gegenfurtner, 2006). Electrophysiological recordings in primates have revealed diversity in the preferred hues of color selective neurons in V1 (Lennie et al., 1990; Hanazawa et al., 2000; Hass and Horwitz, 2013), V2 (Kiper et al., 1997; Solomon and Lennie, 2005), V3 (Gegenfurtner et al., 1997), V4 (Bohon et al., 2016) as well as in downstream color selective areas in the ventral visual pathway (Komatsu et al., 1992; Conway et al., 2007). Color tuning functions of individual color-responsive cells measured electrophysiologically have been found to have a range of half-amplitude bandwidths in hue angle from about 30°-170° in V1 (Wachtler et al., 2003) and 20°-85° in V2 (Kiper et al., 1997). The average bandwidth of color tuning functions tends to decrease through the hierarchy of cortical visual areas (Gegenfurtner, 2003), but large populations of broadly tuned neurons are still found in V4 (Bohon et al., 2016).

The color tuning properties of visual cortical neurons can of course interact with tuning properties to other stimulus features. For instance, many cortical neurons are parallelly tuned to both color and luminance (Johnson et al., 2001; Schluppeck and Engel, 2002), spatial frequency (Lennie et al., 1990) or orientation (Leventhal et al., 1995; Friedman et al., 2003; Garg et al., 2019). One distinction that has been relatively well studied is between so-called single- and double color opponent neurons (Shapley and Hawken, 2011). Double opponent neurons respond to stimuli containing sharp chromatic edges but respond weakly or not at all to full-field or low spatial frequency chromatic stimuli, while single opponent neurons show the opposite pattern (Livingstone and Hubel, 1984; Johnson et al., 2008). Depending on the spatial properties of the chosen stimuli, color tuning results may therefore vary, because different populations of cells may be targeted that are tuned to combinations of color and other stimulus features. In particular, the stimuli with chromatic edges may preferentially activate double opponent neurons, while stimuli without edges may preferentially activate single opponent neurons.

In humans, other than by inference from the results of psychophysical methods, there is one existing study by Chen et al. (2021) that measured cortical color tuning functions using neuroimaging, specifically by measuring steady -state visually-evoked potentials (SSVEPs). SSVEPs occur in response to visual stimuli that are flickering at a precise temporal frequency (Regan, 1966; Norcia et al., 2015), and are typically analyzed in the frequency domain where EEG noise is segregated into many frequency bands, providing high signal to noise for brain responses at the stimulus frequency (Rossion and Boremanse, 2011) and a sensitive index of visual responses in the cortex. Chen et al. (2021) used a method for measuring cortical color tuning functions based on chromatic noise masking, where psychophysical results have shown that the strength of masking increases with the color similarity of the target and mask (Gegenfurtner and Kiper, 1992; D’Zmura and Knoblauch, 1998). They presented central binary chromatic noise targets with surround binary chromatic noise masks, and measured occipital SSVEP amplitudes as a function of the angular difference between the color axes from which target and surround colors were sampled, finding broad color tuning functions centered at both cardinal and intermediate color axes.

In the current study we exploited intermodulation of SSVEPs as an alternative scheme for measuring cortical tuning functions (Regan and Regan, 1987). Intermodulation components in SSVEP occur when there are two or more stimuli flickering at different frequencies presented simultaneously, and are thought to provide an index of the degree to which shared neural resources process the different stimuli (Regan and Regan, 1989; Norcia et al., 2015). We presented flickering checkerboard stimuli where odd checks flickered at one frequency and remained at a fixed chromaticity across trials, and even checks flickered at a different frequency and varied in chromaticity. We measured SSVEP amplitudes at the intermodulation frequency, which we expected to be maximal when odd and even checks have the same chromaticity and are therefore processed by the same color mechanisms, and to decrease with the color difference between odd and even checks. In a series of four experiments, we measured cortical color tuning functions at cardinal and intermediate color axes (Experiment 1), when the stimulus lacked perceptible chromatic edges (Experiment 2), as a function of the size of checkerboard checks (Experiment 3), and around the full hue circle (Experiment 4). Our results reveal that bandwidths of cortical color tuning functions do not differ between cardinal and intermediate axes, implying the action of intermediately tuned color mechanisms; that surprisingly, tuning functions do not depend on the presence of perceptible chromatic edges, implying that they probe color mechanisms with low spatial opponency.

## 2.0 General Methods

Provided in this section are methods common to all four experiments.

### 2.1 Participants

Participants in all experiments had normal color vision assessed using an ‘Anomaloskop’ anomaloscope (Oculus, Wetzlar, Germany; Experiments 1-3) or the Ishihara Plates Test (Experiment 4), and were recruited via email and word-of-mouth. Participants provided written informed consent and were compensated £10 per hour for their time. The study received ethical approval from the Sussex University Science and Technology Cross Schools Research Ethics Committee (ER/DJW41), and adhered to the tenants of the World Medical Association’s Declaration of Helsinki (2013) with the exception that it was not pre-registered.

### 2.2 Equipment

Electroencephalographic data were acquired using 64-channel Waveguard caps (ANT neuro, Hengelo, Netherlands) via a high-speed 64-channel ANT Neuro amplifier sampling at 1000 Hz. EEG data were recorded in ASA lab (ANT Neuro, Hengelo, Netherlands). Additional electrodes were used to measure horizontal (HEOG) and vertical (VEOG) eye-movements to allow for the removal of blink artefacts. A custom photodiode input to the EEG amplifier was fitted to check for accurate stimulus timing (see Supplementary Information 7).

Stimuli were displayed on a GDM FW900 CRT monitor (Sony, Tokyo, Japan). The monitor was gamma-corrected using a luminance meter (Konica-Minolta, Tokyo, Japan) and color calibrated using a PR650 SpectraScan spectroradiometer (PhotoResearch, Dorking, Surrey). Stimuli were presented using a ViSaGe MKII Stimulus Generator (Cambridge Research Systems, Rochester, UK) using Matlab 2017 (The Mathworks, Nantick, CA). Triggers to the EEG amplifier from the ViSaGe via BNC and parallel port connections. The monitor had a resolution of 800 x 600 pixels and ran at a frame rate of 160 Hz.

### 2.3 Heterochromatic flicker photometry

Heterochromatic flicker photometric measurements were made in order to calculate stimuli that are isoluminant for individual observers. Stimuli consisted of a 2D annulus with an outer diameter of 41.9° and inner diameter of 5.7°, which oscillated at 11.4 Hz between two colors (Figure 1). On each trial, the intensity of one color was fixed, and participants adjusted the intensity of the other color until they perceived minimum flicker (the point of isoluminance). On the basis of these measurements, stimuli in the four experiments were made isoluminant to ensure that they isolated chromatic mechanisms and did not stimulate luminance mechanisms.

**Figure 1.**
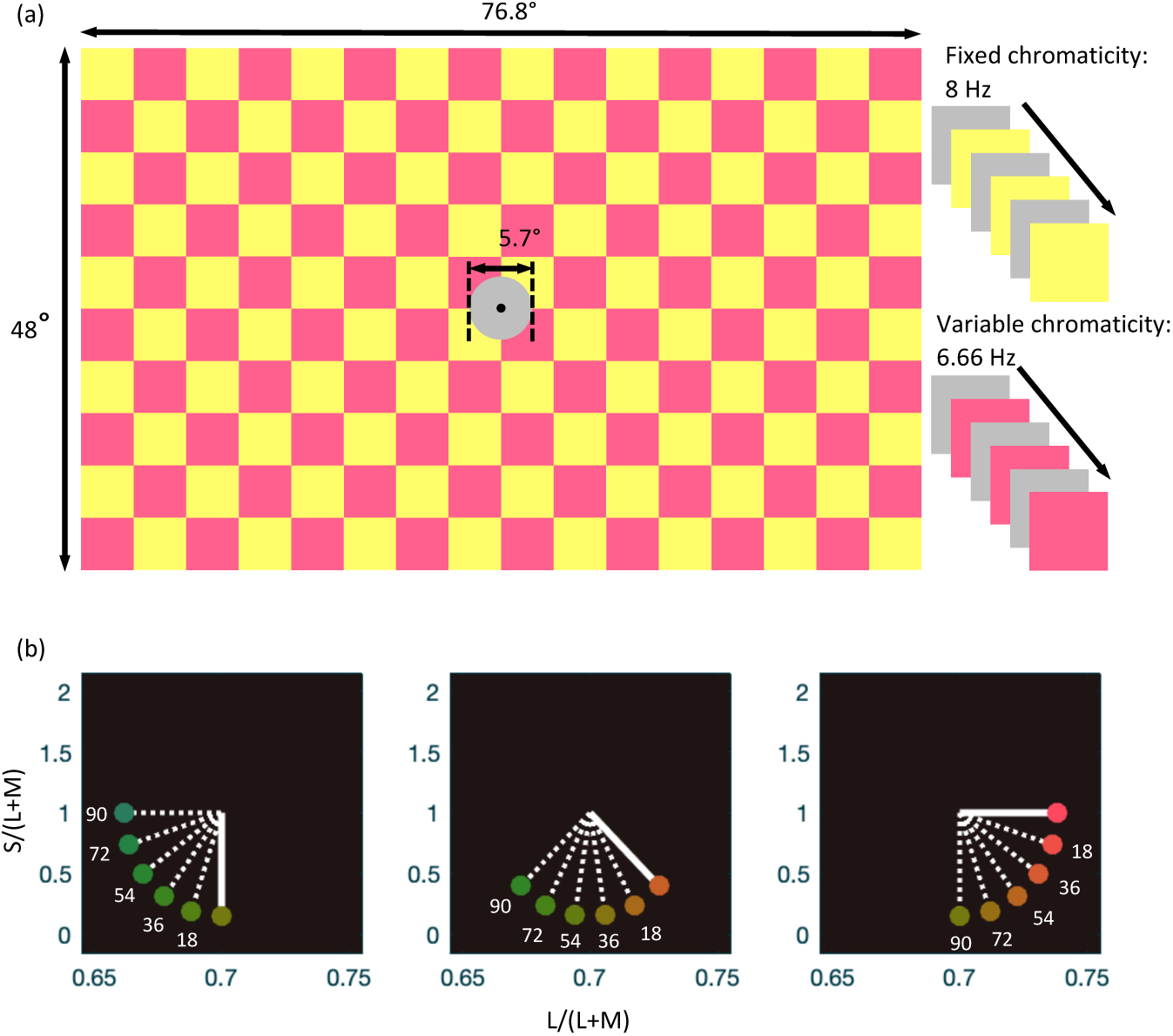
Stimulus properties. (a) Stimuli were isoluminant chromatic flickering checkerboards. Even checks flickered at F_A_ (6.66 Hz) between equal energy white and a ‘variable’ chromaticity that depended on the color condition. Odd checks flickered at F_B_ (8 Hz) between equal energy white and a ‘fixed’ chromaticity. A central 5.7° disc was metameric with equal energy white and did not flicker. (b) Stimulus chromaticities specified in a version of the MacLeod-Boynton (1979) chromaticity diagram based on the Stockman, MacLeod and Johnson (1993) cone fundamentals. For Experiment 1 the fixed chromaticity was either the S/(L+M) axis in a decremental direction (left panel), an intermediate axis (middle panel) or the L/(L+M) axis in an incremental direction (right panel). In each panel, the modulation made by the fixed chromaticity and equal energy white is indicated by the solid line. For each fixed color axis there were 6 color conditions, where the variable chromaticity differed from the fixed chromaticity in steps of 18° from 0° to 90°. Modulations between variable chromaticities and equal energy white are indicated by the dotted lines. For Experiments 2 and 3 the S/(L+M) axis in a decremental direction was tested (left panel). For Experiment 4 the fixed chromaticity was along the S/(L+M) axis in a decremental direction (left panel) and there were 12 color conditions in 30° steps between 0° and 300° from the fixed chromaticity (not shown).

### 2.4 Stimuli

The stimuli for all four experiments consisted of flickering checkerboards extending 76.8° horizontally and 48° vertically (Figure 1a), with a viewing distance of 30 cm. Each check was square and measured 4.8° x 4.8°. Even checks flickered between equal energy white and a fixed hue at 8 Hz, eliciting SSVEPs at 8 Hz and higher harmonics. Odd checks flickered between equal energy white and a variable hue at 6.66 Hz, eliciting SSVEPs at 6.66 Hz and higher harmonics. Flicker was square wave between equal energy white and the edge of the maximum saturation circle allowed by the monitor’s gamut (Figure 1b). There was a central disc metameric with equal energy white of 5.7° with a central fixation point which did not flicker, to avoid the macular pigment (Figure 1a).

### 2.5 Procedure

Participants first completed heterochromatic flicker photometry task. The consistency of intensity settings was assessed, and if found to range >20%, the task was repeated. Participants were then fitted with facial electrodes and an EEG cap in accordance with the 10/20 system (Jurcak et al., 2007). We ensured that impedances for electrodes of interest were < 10 kΩ and for other electrodes < 20kΩ. In the experiment, participants viewed a series of 10 second trials (see details of methods for each separate experiment). They were instructed not to blink excessively during trials and to blink between trials. Participants controlled the inter-trial interval by pressing a button on a keyboard when they were ready for the next trial. Raw EEG data were monitored in real time for signs of fatigue or other issues.

### 2.6 EEG Analysis: pre-processing

Pre-processing was conducted in MATLAB, using Letswave 7 (https://nocions.github.io/letswave7/) for Experiments 1 and 4, and Letswave 6 for Experiments 2 and 3 (https://nocions.github.io/letswave6/). First, raw EEG data were imported and mapped to the ANT neuro standard 64-channel electrode coordinate system. Raw data were visually inspected for excessive noise resulting from equipment failures. Problematic channels were removed. No exclusions of channels were made from our analysis cluster, consisting of Oz, O1, O2, Pz, P1, P2, P3, P4, POz, PO3, and PO4 (Figure 2d). Overview of tuning functions at other posterior and occipital channels is presented in Supplementary Information 5 (Supplementary Figure 6), demonstrating signal transitivity across the area. Signals were re-referenced to the grand average of all remaining cortical electrodes. In addition to our analysis cluster, channels FPz, HEOG and VEOG were extracted to detect eye movements. Consistency in temporal profile of our stimuli was ensured by inspecting the decomposed frequency profile of the photodiode which was attached to the screen (see Supplementary Information 7). A Butterworth filter for 50 Hz noise and a high-pass filter (<0.1 Hz) were applied. Data were then segmented into trials using an epoch size of 9000 s, which included an integer number of stimulus repetitions at both flicker frequencies. Segmented trials were baseline corrected within the 1 to 1.2 interval. No ocular artefacts were removed in Experiments 1 and 4 (see Supplementary Information 2 for a comparison of signal amplitudes with and without ocular artefact removal). For Experiments 2 and 3, ocular artefacts were removed by visual inspection if there were fewer than 2 blinks per color condition. Alternatively, Independent Components Analysis (ICA) was applied if there were two or more blinks per condition. Recordings for an individual observer were merged in cases where data was collected across multiple sessions. Trials from the same condition for each observer were then averaged and converted to the frequency domain by fast Fourier transformation (FFT). Amplitudes for each observer and each color condition were extracted at I_1_ (14.66 Hz), which is an intermodulation component at the sum of the two stimulus frequencies. Amplitudes were also extracted at F_B_ (8 Hz), the frequency of the checks of fixed chromaticity – changes in amplitude with the chromaticity of the F_A_ (6.66 Hz) checks reflects chromatic masking, which has previously been used to infer cortical color tuning functions (Chen and Gegenfurtner, 2021).

**Figure 2.**
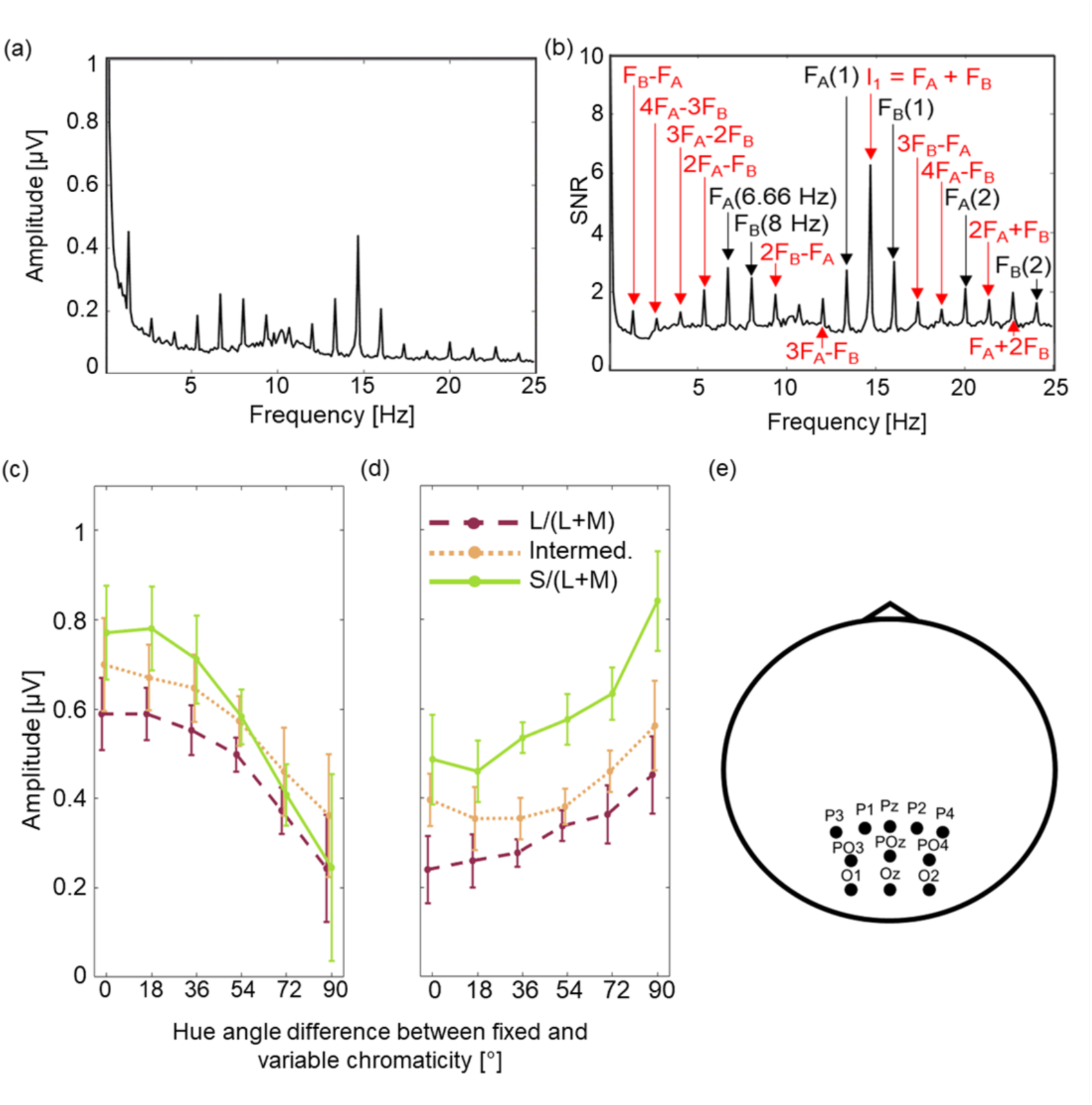
Experiment 1 results. (a) Amplitude profile for an example condition (S/(L+M) axis, 0° variable chromaticity), averaged over the 11 participants. (b) SNR profile for the same condition. Fundamental stimulus frequencies and their harmonics are labelled in black and intermodulation components in red. (c) Mean amplitude of the intermodulation component at 14.66 Hz as a function of the hue angle between the fixed and variable chromaticities for the S/(L+M) axis (green solid line), the intermediate axis (orange dotted line) and the L/(L+M) axis (red dashed line). Error bars are 95% confidence intervals. (d) Mean amplitude at the stimulus frequency of the fixed chromaticity at 8 Hz as a function of the hue angle between the fixed and variable chromaticities for the S/(L+M) axis (green solid line), the intermediate axis (orange dotted line) and the L/(L+M) axis (red dashed line). Error bars are within-participant 95% confidence intervals (O’Brien and Cousineau, 2015), jittered along the x-axis for visibility. (e) Schematic representation of analysis cluster electrodes.

## 3.0 Experiment 1

In Experiment 1 we sought to measure cortical color tuning functions using SSVEP, and to determine whether there are differences in cortical color tuning functions between ‘cardinal’ color axes aligned with the retinogeniculate color channels L/(L+M) and S/(L+M) and an intermediate color axis.

### 3.1 Experiment 1: Methods

Eleven participants (4 female) completed two experimental sessions, each lasting 2.5 hours. Three color axes were tested: L/(L+M) increments, S/(L+M) decrements and an intermediate axis (Figure 1b). For each axis, the chromaticity of checks flickering at F_A_ was fixed along that axis, and the chromaticity of checks flickering at F_B_ varied on each trial between six hue angles to the fixed axis (0°, 18°, 36°, 54°, 72° and 90°; Figure 2a). There were 180 trials in each session presented in a random order, totaling 360 trials.

### 3.2 Experiment 1: Results

Strong signals were observed at the two fundamental frequencies, their harmonics (F_A_(1) and F_B_(1)), and at I_1_ (e.g. Figures 2a and b). There were additional amplitude peaks at other harmonics of F_A_ and F_B_ and at other intermodulation frequencies. We extracted I_1_ and F_B_ for further analyses.

Figure 2 shows the effect of hue dissimilarity between odd and even checks on SSVEP amplitudes at I_1_ (panel c) and F_B_ (panel d) for the three color axes. Tuning functions were similar in shape for all three axes, with a broad half-amplitude bandwidth of approximately 90°. Two-way repeated-measures ANOVAs were conducted to test the effect of color axis and hue angle differences on the amplitudes of I_1_ and F_B_. Mauchly’s test of sphericity was violated for both frequencies (*X*^2^(54) = 106.50 for I_1_ and *X*^2^ (54) = 115.40 for F_B_, *p* < .001), therefore Greenhouse-Geisser corrected tests were used (ɛ =.311 for I_1_ and .260 for F_B_). The ANOVAs showed that for both I_1_ and F_B_ there were significant main effects of hue angle difference (*F*(1.19,11.92) = 21.59 for I_1_ and *F*(1.84,15.49) = 23.18 for F_B_, *p* <.001, ⴄ^2^_p_ =.683 for I_1_ and ⴄ^2^_p_ = .699 for F_B_). Results from post-hoc tests (Supplementary Information 1, Supplementary Tables 1 and 2) show that mean SSVEP amplitude decreased significantly with hue angle difference for I_1_ and increased significantly with hue angle difference for F_B_.

There was a significant main effect of color axis for F_B_, (*F*(1.14, 11.36) = 10.83, *p* = .006, ⴄ^2^_p_ =.520), but not for I_1_ (*F*(1.45,14.51) =3.17, *p* =.084, ⴄ^2^p =.241). Post-hoc tests using Bonferroni correction indicated that there was a significant difference between L/(L+M) and S/(L+M) axes (*M* = -.247, *p* < .05). There was no significant interaction between axis and hue angle difference either for I_1_ (*F*(3.11,31.06) = 2.74, *p* = .058) nor for F_B_ (*F*(2.60, 26.03) = 2.40, *p* = .098). Thus, the effect of hue angle on SSVEP amplitude was consistent across all three axes for both frequencies, implying that tuning functions were shaped similarly for all three axes.

## 4.0 Experiment 2

The isoluminant checkerboard stimuli in Experiment 1 have strong chromatic edges. The S/(L+M) axis, and thus neural populations responding to chromatic edges may underlie the results. To test the effect of chromatic edges on cortical color tuning functions measured using SSVEP we reduced the size of the ‘checks’ to single pixels (invisible to participants). If tuning functions change from those measured in Experiment 1 when the checkerboard is constructed from single pixels, then the tuning functions measured in Experiment 1 are attributable to neural populations responsive to chromatic edges.

### 4.1 Experiment 2: Methods

Eleven participants (6 female) completed a single 2.5-hour experimental session. Stimuli consisted of single pixel checkerboards (.096° x .096°), with the same spatial organization as depicted in Figure 1a. Individual checks were indistinguishable to the observer. The chromaticity of checks flickering at F_B_ was fixed, and the chromaticity of checks flickering at F_A_ varied between six hue angles to the fixed chromaticity (0°, 18°, 36°, 54°, 72° and 90°). There were 120 experimental trials, where hue conditions were presented in a random order. Since the S/(L+M) axis produced somewhat larger amplitudes in Experiment 1 than the other two axes, Experiment 2 probed the S/(L+M) axis only.

### 4.2 Experiment 2: Results

Figure 3 shows the amplitudes of I_1_ and F_B_ against the difference in hue angle between odd and even single pixel checks. One-way repeated-measures ANOVAs examined the effect of hue angle difference on the amplitudes of I_1_ and F_B_. Mauchly’s test of sphericity was violated for both frequencies (*X*^2^(14) = 72.16 for I_1_ and *X*^2^(14) = 34.53 for F_B_, *p* <.001), therefore Greenhouse-Geisser corrected tests were used (ɛ = .279 for I_1_ and ɛ = .510 for F_B_). There was a significant effect of hue angle difference on signal amplitude for I_1_ (*F*(1.40, 13.96) = 9.56, *p* =.005, ⴄ^2^p = .489), but not for F_B_ (*F*(2.55, 25.48) = .465, *p* = .679, ⴄ^2^_p_ =.044). Results from post-hoc tests (see Supplementary Information 1, Supplementary Tables 3 and 4) indicated that mean SSVEP amplitude decreased with hue angle difference for I_1_, but did not change with hue angle difference for F_B_.

**Figure 3.**
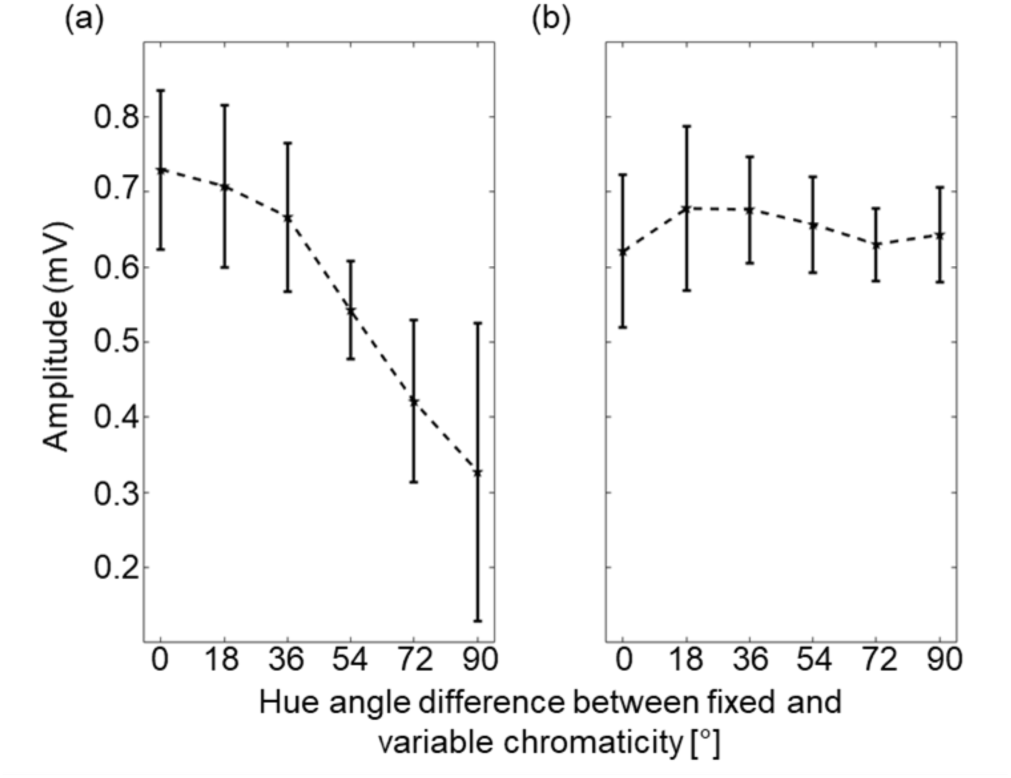
Results of Experiment 2. SSVEP amplitude is plotted as a function of hue angle dissimilarity for single-pixel checkerboard stimuli. (a) for I_1_ at 14.66 Hz. (a) for F_B_ at 8 Hz. Error bars are within-participant 95% confidence intervals (O’Brien and Cousineau, 2015).

## 5.0 Experiment 3

Since we observed a difference in the shape of the tuning function for F_B_ between the 4.8° checks presented in Experiment 1 and the indivisible 0.096° checks presented in Experiment 2, in Experiment 3 we systematically manipulated check size. We measured tuning functions at I_1_ and at F_B_ at 4 check sizes, varying between the check sizes used in Experiments 1 and 2.

### 5.1 Experiment 3: Methods

Eleven participants (5 female) completed two 2.5-hour experimental sessions. There were 4 check size conditions: 0.096° (as in Experiment 2), 0.4°, 1.3° and 4.8° (as in Experiment 1). To limit the duration of testing sessions there were four hue angle difference conditions rather than the 6 used in Experiments 1 and 2. These were 0°, 30°, 60° and 90°. Checks flickering at F_B_ (8 Hz) at a fixed chromaticity modulated between equal energy white and a decrement along S/(L+M) axis (left panel of Figure 1b). 2 testing sessions were conducted separated by at least one day. There were 160 experimental trials per session (10 trials for each combination of check size and hue angle difference), totaling 320 trials (20 trials in total for each stimulus type).

### 5.2 Experiment 3: Results

Figure 4 shows the effect of check size and hue angle difference on amplitudes at I_1_ and F_B_. The figure shows that for I_1_ check size does not affect the shape of the tuning functions observed. However, F_B_, in agreement with the results of Experiment 2 there is no tuning for the single pixel checkerboard, but similar tuning functions as observed in the results of Experiment 1 for the larger check sizes.

**Figure 4.**
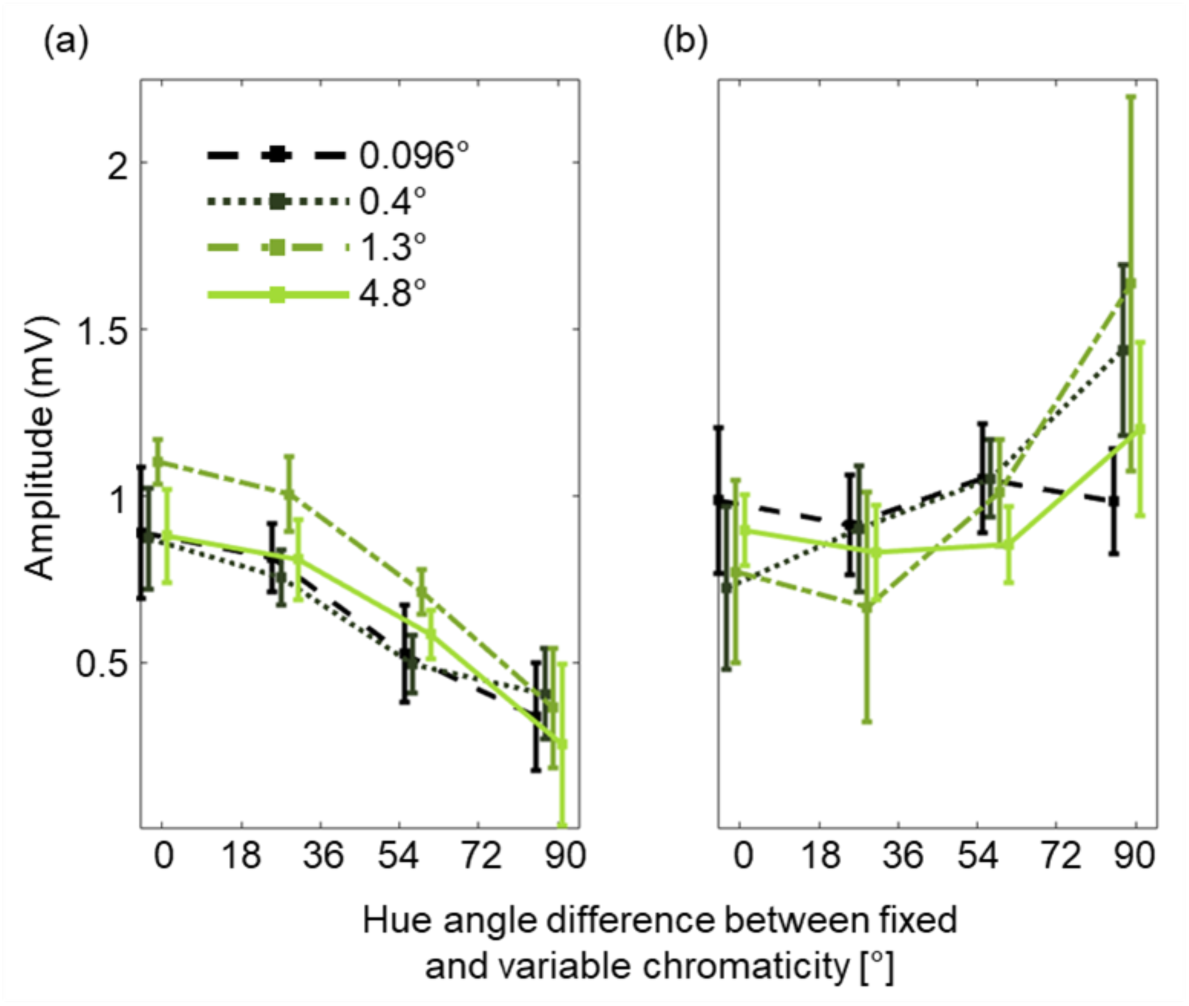
Results of Experiment 3. SSVEP amplitude is plotted as a function of hue angle dissimilarity (x-axis) and check size (legend). (a) for I_1_ at 14.66 Hz. (b) for F_B_ at 8 Hz. Error bars are within-participant 95% confidence intervals (O’Brien and Cousineau, 2015), jittered along the x-axis for visibility.

We conducted two-way repeated-measures ANOVAs to test the effect of hue angle hue angle difference and check size on the amplitudes at I_1_ and F_B_. Mauchly’s test of sphericity was violated for both frequencies (*X*^2^(44) = 81.52, *p* = .002 for I_1_; *X*^2^(44) = 134.05, *p* < .001 for F_B_). Greenhouse-Geisser corrected tests were therefore used (ɛ = .367 for I_1_; ɛ = .251 for F_B_). There were significant main effects of check size and hue angle on signal amplitude for I_1_ (*F*(2.32, 23.21) = 3.61, *p*= .038, ⴄ^2^_p_ = .265 for check size; *F*(1.24, 12.39) = 33.76, *p* <.001, ⴄ^2^_p_ =.771 for hue angle difference), but no significant interaction (*F*(3.30, 33.01) = 1.55, *p* =.218, ⴄ^2^_p_ =.134). Post-hoc tests (Supplementary Information 1, Supplementary Tables 5 and 6) indicated that mean amplitude in Condition 3 was somewhat stronger than in other conditions, regardless of hue angle, and that mean SSVEP amplitude decreased with hue angle for I_1_, regardless of check size. For F_B_ there was a significant main effect of hue angle difference (*F*(1.27, 12.67) = 11.30, *p* = .004, ⴄ^2^_p_ =.530), but there was no significant main effect of check size (*F*(1.51, 15.06) = .428, *p* =.604, ⴄ^2^_p_ =.041). Results from post-hoc tests (Supplementary Information 1, Supplementary Tables 5 and 6) indicated that mean SSVEP amplitude increased with hue angle difference. There was a significant interaction between check size and hue angle for F_B_ (*F*(2.26, 22.58) = 3.39, *p* = .046, ⴄ^2^_p_ =.253). Post-hoc analysis (Supplementary Information 1, Supplementary Table 6) indicated that the shape of the tuning function for F_B_ differed between check size conditions.

## 6.0 Experiment 4

The tuning functions observed in the results of Experiments 1-3 are broadband but do not reach an asymptote at the largest hue angle differences we measured of 90°. In Experiment 4 we aimed to characterize a tuning function for a cortical color mechanism around the entire hue circle. We kept the fixed chromaticity checks along the S/(L+M) axis in a decremental direction but varied the variable chromaticity checks around the full hue circle in 12 steps.

### 6.1 Experiment 4: Methods

Ten participants (7 female) completed Experiment 4 in one experimental session lasting about 3 hours. Stimuli were isoluminant flickering checkerboards with a check size of 1.3° (which elicited the largest signals in Experiment 3). The fixed chromaticity flickering at F_B_ (8 Hz) was a decrement on the S/(L+M) axis (first panel of Figure 1b). The chromaticity of the variable checks flickering at F_A_ (6.66 Hz) spanned the full hue circle in steps of 30° (hue angle differences ranged from 0° to 330°). Each of the 12 hue angle conditions was presented 20 times, totaling 240 trials. SSVEP amplitudes were measured at I_1_ and at F_B_.

### 6.2 Experiment 4: Results

Results from Experiment 4 are presented in Figure 5a. There is a peak in I_1_ centered at 0° hue angle difference, as expected from the results of Experiments 1-3. However, there is a second smaller peak centered at a hue angle difference of 180°. For F_B_ there is a minimum at a hue angle difference of 0° as expected from the results of Experiments 1 and 3. There is no evidence of a second minimum at a hue angle difference of 180°.

**Figure 5.**
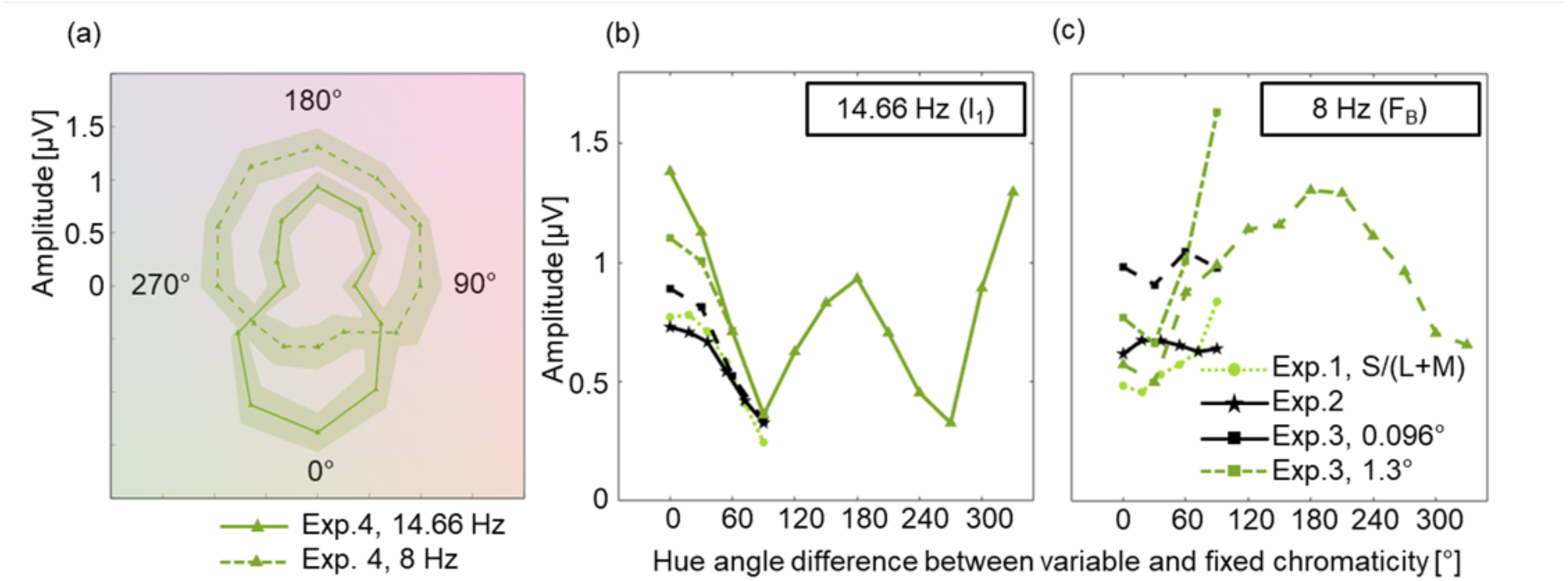
Results of Experiment 4. (a) tuning functions plotted at I_1_ (full line) and F_B_ (dashed line), where the fixed chromaticity was an S/(L+M) decrement (180°), and the variable chromaticity varied around the full hue circle. Error envelopes are within-participant 95% confidence intervals (O’Brien and Cousineau, 2015). Tuning functions for decrement in S/(L+M) mechanism comparing Experiment 4 results with those from Experiments 1-3 are plotted in Panel (b) for I_1_ (14.66 Hz) and Panel (c) for F_B_ (8 Hz). For Experiment 3, results are shown only for the 0.096° and 1.3° check size condition.

One-way repeated measures ANOVAs were used to investigate the amplitude differences between hue angle difference conditions at I_1_ and at F_B_. There was a significant effect of hue angle difference on amplitudes at I_1_ (*F*(11, 99) = 18.065, *p* <.001, ⴄ^2^_p_ = .667) and at F_B_ (*F*(11, 99) = 9.851, *p* <.001, ⴄ^2^_p_ = .523). Post hoc comparisons (Supplementary Information 1, Supplementary Tables 7 and 8) showed that amplitudes at I_1_ for a 0° hue angle difference were significantly higher than amplitudes at all other angles, except for 330°. The amplitude for a hue angle difference of 180° was not significantly different from those of its most proximal hue angle difference conditions, 150° and 210°.

## 7.0 Analyses of additional intermodulation components

It is thought that different intermodulation components may arise from separate subpopulations of neurons (Regan, 1966; Appelbaum et al., 2008; Gordon et al., 2019). For example, the subpopulation of neurons responding to the difference between F_A_ and F_B_ might be separate from the population responding to the sum of F_A_ and F_B_, even though they both arise from interactions to the same signals (F_A_ and F_B_). These potentially different populations may also demonstrate distinct color tuning and may be differently spatially distributed across the cortex. We therefore conducted additional analyses of the data from Experiments 1 and 4 to plot tuning functions for other intermodulation components and visualize their locations in cortical electrode maps. Firstly, we extracted SNR profiles for a grand mean across observers in the 0° hue difference condition (where intermodulation components are expected to be greatest). Intermodulation components (i.e., those where the frequency peak is composed of either an addition or subtraction of multiples of F_A_ and F_B_) were extracted when the SNR peak surpassed a criterion of 2. This procedure identified 2 qualifying components in Experiment 1 (Supplementary Information 4, Supplementary Figure 5) and 12 qualifying components in Experiment 4 (Figure 6). One response at 4 Hz (labelled orange in Figure 6) could either be an intermodulation component (3F_B_ - 3F_A_) or a subharmonic of F_B_ (F_B_/2). Since we found no peak at the equivalent subharmonic of F_A_, this peak is more likely to be an intermodulation component. Tuning functions were calculated for the channels in the analysis cluster and visualized using scaled line plots. Tuning functions for different intermodulation components had different shapes (e.g., compare tuning functions for F_A_+F_B_ and for 2F_A_+2F_B_). However, since the criterion SNR was applied to the group average, data from some participants had high noise, impacting the group average tuning functions.

**Figure 6.**
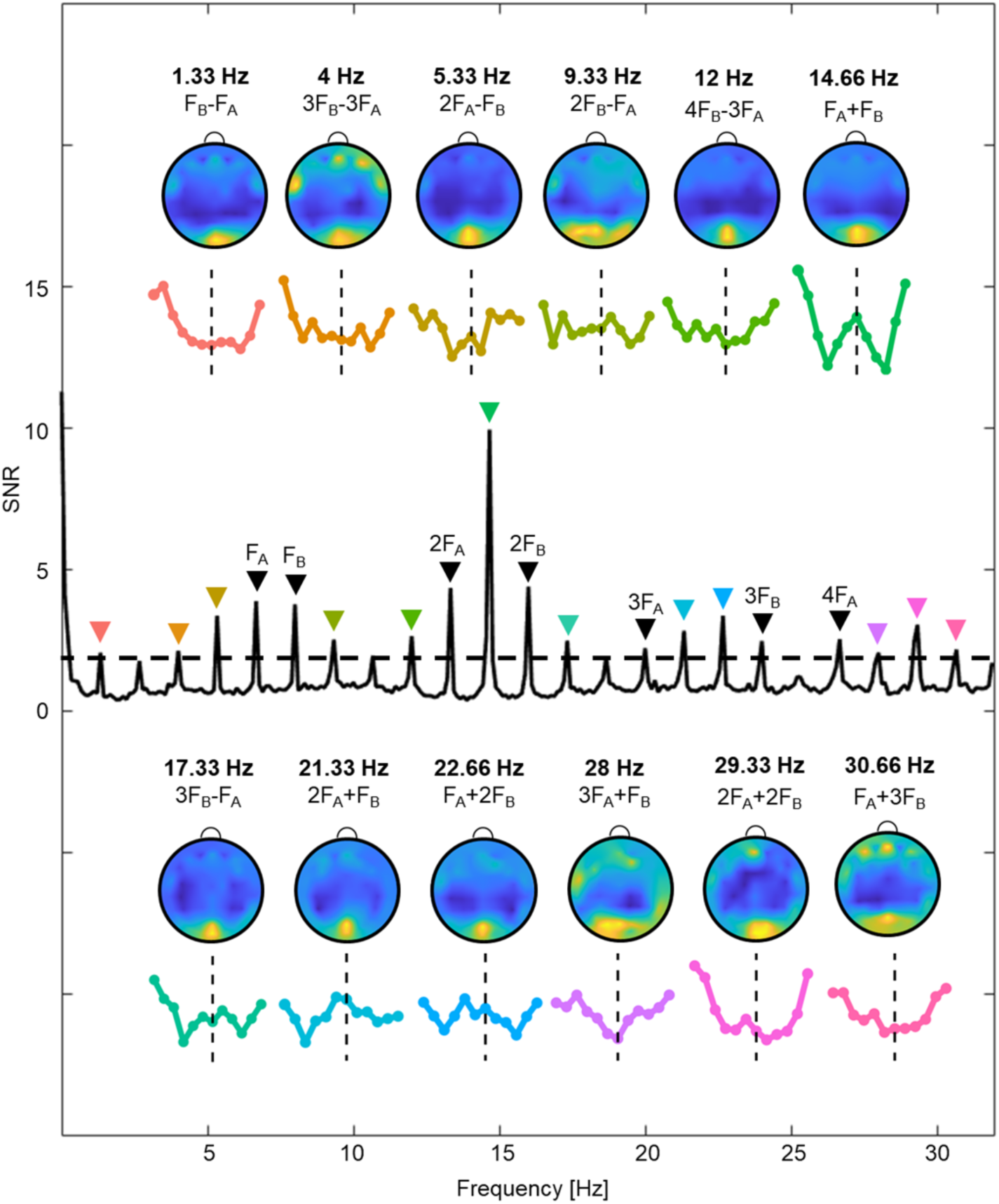
Signal heatmaps and tuning functions at intermodulation components surpassing 2 SNR in Experiment 4. SNR is presented as a function of Frequency in Hz with a range of 0 – 25. The components and their sources (nature of interaction between F_A_ and F_B_) are labelled above individual heatmap and tuning function plots. The individual heatmaps and tuning function plots are scaled to match in area under the curve.

Spatial signal distributions were visualized using amplitude scalp heatmaps with the grand mean across observers, using a relativized scale between frequencies so that the maximum values are always plotted in yellow and minimum values in dark blue. Comparison of the cortical surface heatmaps for different intermodulation components revealed that their locations are broadly consistent, both for Experiment 1 and Experiment 4. However, we found some substantial differences in tuning functions for different intermodulation components. In results for Experiment 4, Some functions show a *reduction* in amplitude for the opponent hue (e.g., 26.66 Hz and 28 Hz), in contrast to the characteristic peak observed in I_1_. At the 20 Hz intermodulation component, tuning appears to be reversed, with a higher relative peak for the opponent compared to the proximal hues (Figure 6). Overall, peaks are less pronounced for intermodulation components other than I_1_, but nonetheless may reveal different color tuning properties for different neural subpopulations.

## 8.0 Discussion

Our results show that human cortical color tuning functions can be effectively measured by capturing intermodulation of SSVEPs in EEG. We separately frequency tagged two colors spatially interleaved in checkerboard patterns and extracted signals at the intermodulation frequency (I_1_). As predicted, SSVEP amplitudes at I_1_ significantly and systematically reduced with hue angle dissimilarity. All of the tuning functions we measured by extracting I_1_ were broad, with half-amplitude bandwidths of about 90° in hue angle. This bandwidth is concordant with existing reported bandwidths of color tuning functions measured psychophysically (Sankeralli and Mullen, 1997; Lindsey and Brown, 2004) and electrophysiologically (Wachtler et al., 2003). The results of Experiment 1 showed similar tuning functions for cardinal and intermediate color axes. The results of Experiments 2 and 3 showed that the shape of color tuning functions did not depend on whether or not the stimulus contained perceptible edges or the sizes of checkerboard checks in the stimulus. The results of Experiment 4 showed that tuning functions around the whole hue circle exhibit a bipolar shape with similar 90° half-amplitude bandwidths at both peaks. We interpret each of these findings in turn.

To assess the independence or interaction between color representations along ‘cardinal’ retinogeniculate color axes and intermediate color axes, in Experiment 1 we measured tuning functions centered on both cardinal axes and on an intermediate axis. If colored stimuli specified along an intermediate axis are represented by a combination of activities in the two cardinal mechanisms, then we would expect observed ‘tuning functions’ along the intermediate axis to have a larger bandwidth and a smaller peak amplitude than tuning functions for stimuli with equivalent contrast along the cardinal axes. However, in the results of Experiment 1 we found very similar peak amplitudes and bandwidths for tuning functions centered along an intermediate axis as for the cardinal axes, implying the action of an independent intermediately tuned cortical color mechanism. How are the cardinal retinogeniculate color mechanisms combined in the cortex to produce intermediate color mechanisms? Gegenfurtner and colleagues have suggested that broad tuning with a half-amplitude bandwidth greater than 30° typically results from a linear combination of cone-opponent signals (Hansen and Gegenfurtner, 2006; Chen and Gegenfurtner, 2021). However, nonlinear combinations of cone-opponent signals could also result in broad tuning functions along intermediate axes depending on how the inputs are combined. In their ‘hue sweep’ SSVEP study, Kaneko et al. (2020) found that SSVEP amplitudes in response to colors along intermediate axes could be greater than SSVEP amplitudes in response to colors along the cardinal axes, which they interpreted as strong evidence that intermediately tuned color mechanism result from nonlinear combinations of the cardinal mechanisms. In our Experiment 1 results, the SSVEP amplitudes of I_1_ along the intermediate axis were approximately midway between those of the two cardinal axes. This pattern of results, according to Kaneko et al. is in favor of a nonlinear combination of cardinal mechanisms. But the results of both studies could also be explained by linear combination followed by independent gain control for each second stage color mechanism. A linear combination of cardinal mechanisms to form intermediate mechanisms would be in accordance with recent optical imaging evidence for linear combination in macaque V1 (Li et al., 2022).

To differentiate the contributions of single- and double-opponent neurons to the cortical color tuning functions that we measured in Experiment 1, in Experiments 2 and 3 we manipulated chromatic edges in the stimulus. In Experiment 2, our ‘single pixel’ checkerboard stimulus had no perceptible chromatic edges and should therefore excite only single-opponent neurons. We found, surprisingly, that tuning functions based on I_1_ had the same shape and bandwidth as the tuning function based on the large checks with chromatic edges in Experiment 1. In Experiment 3 we manipulated check size systematically over 4 levels, and found no effect on the bandwidth of I_1_ with check size. We found that check sizes of 1.3° generated slightly higher amplitudes than the other check sizes. However, the lack of significant interaction between check size and hue angle dissimilarity suggests that the larger amplitudes could result from responses by other spatial-frequency tuned neural populations that are not hue-angle specific. As well as SSVEP amplitudes at I_1_ we also extracted amplitudes at F_B_, the frequency (8 Hz) at which the ‘fixed chromaticity’ was modulated. Chen and Gegenfurtner (2021) had previously plotted cortical color tuning functions using this frequency, reasoning that amplitudes reduce with chromatic masking if the underlying color channel is sensitive to both modulating colors. We found, in contrast to results at our main target I_1_, that tuning functions plotted using F_B_ did depend on chromatic edges. For the larger checks used in Experiments 1 and 3 (similar to those used by Chen and Gegenfurtner), tuning functions derived from amplitudes at the frequency of the fixed chromaticity F_B_ were roughly the inverse of tuning functions derived from I_1_, implying that both reveal the action of the same chromatic mechanisms. However, in Experiments 2 and 3, the shapes of the observed tuning functions derived from F_B_ differed substantially from those derived from I_1_, suggesting that they arise from distinct chromatic mechanisms. For our single pixel checkerboard stimuli, tuning functions derived from I_1_ matched those measured using larger check sizes, but tuning functions derived from F_B_ were flat. The mechanism revealed by F_B_ is therefore dependent on spatial chromatic edges, but the mechanism revealed by I_1_ is not. The neural populations underlying F_B_-derived tuning functions may be double-opponent while those underlying I_1_-derived tuning functions may be single-opponent.

In Experiments 1-3 the chromaticity of the “variable” checks was manipulated between hue angle differences of 0° and 90°. To measure the shape of a color tuning function around the full hue circle, in Experiment 4 we manipulated the chromaticity of the variable checks between hue angle differences of 0° and 360°. Surprisingly, we found a second peak in I_1_ at a hue angle difference of 180°, where the hue of the variable chromaticity was opponent to the hue of the fixed chromaticity. The second peak had about half the amplitude of the primary peak centered at a hue angle difference of 0°. We did not observe this pattern for tuning functions based on F_B_, where we observed a single minimum at 0° and a single maximum at 180°. What could explain this pattern of findings? We assume, as argued by Chen and Gegenfurtner (2021) that tuning functions plotted by extracting F_B_ are based on chromatic masking. Is chromatic masking more effective for colors 90° distinct from the target than for colors 180° distinct from the target? We have not identified any psychophysical data that could address this hypothesis, but it seems unlikely that this would be the case: at 90° masking should be minimal as mask and target chromaticities are carried by orthogonally tuned color mechanisms. However, target signals will also be influenced by simultaneous contrast, where a surround chromaticity pushes perception of a target towards the surround’s complementary color (Ekroll and Faul, 2012). Simultaneous color contrast induced by surround S-cone increment checks would enhance color signals from S-cone increment checks, but simultaneous color contrast induced by L/(L+M) increment or decrement checks would not affect color signals from S-cone increment checks as it would act orthogonally to the S-cone axis. The single peak in tuning for F_B_ could therefore be explained by a combination of masking and simultaneous contrast.

For tuning functions plotted by extracting I_1_, the second peak in amplitude at a hue angle difference of 180° must arise when the same neural populations are driven by both checks. The checks with fixed color modulated in chromaticity between an S-cone decrement and equal energy white. The variable color checks at a 180° hue angle difference modulated between equal energy white and an S-cone increment. One possibility is that a color mechanism with bipolar tuning (Conway et al., 2002; Chatterjee and Callaway, 2003) responds to both sets of checks, leading to the second intermodulation peak observed at a hue angle difference of 180°. A second possibility is that the intermodulation could arise within rectified S-cone on and off channels, which would be consistent with multiple unipolar channels for color in the cortex (Malkoc et al., 2005; Emery et al., 2017, 2023). S-cone off channels would respond to the onset of S-cone decrement checks (at F_A_) and to the offset of S-cone increment checks (at F_B_). S-cone on channels would show the opposite pattern of responses. Intermodulations could thus occur separately within on and off channels but these separate responses would both be at the same intermodulation frequency and would not be separable in our results. Further work could attempt to measure responses in S-cone increment and decrement channels separately by manipulating the temporal waveform (Krauskopf and Zaidi, 1986; Stockman et al., 2017).

SSVEP responses as captured by EEG reflect activity from multiple neural populations. However, it may be possible to capture the activity of different neural populations by investigating different intermodulation components (Regan, 1966; Appelbaum et al., 2008; Alp et al., 2016; Gordon et al., 2019). We therefore plotted color tuning functions for all the intermodulation components that met our criterion of 2 SNR in Experiments 1 and 4. By eye, we can see two distinct types of tuning function in the results of Experiment 4 (Figure 6). Tuning functions based on some of the intermodulation components show a second peak at a 180° hue angle difference as for I_1_ (3F_B_-F_A_, 2F_A_+F_B_ and F_A_+2F_B_), while tuning functions based on others do not have the second peak at a 180° hue angle difference (F_B_-F_A_, 3F_B_-3F_A_, 6F_B_-3F_A_, 2F_A_+2F_B_ and F_A_+3F_B_). It is possible that these different groups of intermodulation components reflect the actions of different neural populations with different color tuning properties. However, the scalp map distributions of the different components are not obviously different (Figure 6). We also did not find any categorical effect of difference intermodulation components versus sum intermodulation as has been suggested in the literature (e.g., the tuning functions for F_A_+F_B_ and 2F_A_+2F_B_ are both sum intermodulation components but have different shapes).

Our investigation of intermodulation of EEG signals has revealed broadband cortical color tuning functions. We found that cortical tuning is consistent across both cardinal and intermediate color axes, implying the action of intermediately tuned cortical color mechanisms. Surprisingly, the tuning functions we measured with intermodulation of EEG do not rely on chromatic edges, suggesting that they arise from single-opponent neural populations. Our tuning functions are consistent with those derived psychophysically, from single cell electrophysiology and from color masking in EEG, showing that intermodulation SSVEP is a promising method for probing cortical color representation in humans.

## Acknowledgements

The work was funded by ERC grant 949242 COLOURCODE awarded to JMB.

