## Supplementary information for "Tuning of cortical color mechanism revealed using steady-state visually evoked potentials"

#### 1. Post hoc analyses

<u>Experiment 1</u>

**Supplementary Table 1**

*Post hoc tests of interaction between axis and hue angle for $I_1$ (14.66 Hz) in Experiment 1*

| | | Mean Difference | SE | t | Cohen's d | $p_{bonf}$ |
|---|---|---|---|---|---|---|
| L/(L+M), 0° | Intermed., 0° | -0.110 | 0.060 | -1.836 | -0.335 | 1.000 |
| | S/(L+M), 0° | -0.182 | 0.060 | -3.022 | -0.552 | 0.605 |
| | L/(L+M), 18° | < -0.001 | 0.060 | -0.004 | < -0.001 | 1.000 |
| | Intermed., 18° | -0.081 | 0.074 | -1.095 | -0.246 | 1.000 |
| | S/(L+M), 18° | -0.191 | 0.074 | -2.576 | -0.580 | 1.000 |
| | L/(L+M), 36° | 0.037 | 0.060 | 0.611 | 0.111 | 1.000 |
| | Intermed., 36° | -0.058 | 0.074 | -0.782 | -0.176 | 1.000 |
| | S/(L+M), 36° | -0.122 | 0.074 | -1.646 | -0.371 | 1.000 |
| | L/(L+M), 54° | 0.091 | 0.060 | 1.517 | 0.276 | 1.000 |
| | Intermed., 54° | 0.016 | 0.074 | 0.210 | 0.047 | 1.000 |
| | S/(L+M), 54° | 0.006 | 0.074 | 0.084 | 0.019 | 1.000 |
| | L/(L+M), 72° | 0.216 | 0.060 | 3.610 | 0.656 | 0.075 |
| | Intermed., 72° | 0.129 | 0.074 | 1.738 | 0.391 | 1.000 |
| | S/(L+M), 72° | 0.181 | 0.074 | 2.444 | 0.550 | 1.000 |
| | L/(L+M), 90° | 0.346 | 0.060 | 5.773 | 1.049 | < .001 |
| | Intermed., 90° | 0.227 | 0.074 | 3.065 | 0.690 | 0.451 |
| | S/(L+M), 90° | 0.344 | 0.074 | 4.636 | 1.044 | 0.002 |
| Intermed., 0° | S/(L+M), 0° | -0.071 | 0.060 | -1.186 | -0.217 | 1.000 |
| | L/(L+M), 18° | 0.110 | 0.074 | 1.485 | 0.334 | 1.000 |
| | Intermed., 18° | 0.029 | 0.060 | 0.487 | 0.089 | 1.000 |
| | S/(L+M), 18° | -0.081 | 0.074 | -1.087 | -0.245 | 1.000 |
| | L/(L+M), 36° | 0.147 | 0.074 | 1.981 | 0.446 | 1.000 |
| | Intermed., 36° | 0.052 | 0.060 | 0.875 | 0.159 | 1.000 |
| | S/(L+M), 36° | -0.012 | 0.074 | -0.158 | -0.035 | 1.000 |
| | L/(L+M), 54° | 0.201 | 0.074 | 2.712 | 0.611 | 1.000 |
| | Intermed., 54° | 0.126 | 0.060 | 2.104 | 0.382 | 1.000 |
| | S/(L+M), 54° | 0.117 | 0.074 | 1.573 | 0.354 | 1.000 |
| | L/(L+M), 72° | 0.326 | 0.074 | 4.402 | 0.991 | 0.005 |
| | Intermed., 72° | 0.239 | 0.060 | 3.997 | 0.726 | 0.019 |
| | S/(L+M), 72° | 0.292 | 0.074 | 3.932 | 0.885 | 0.027 |
| | L/(L+M), 90° | 0.456 | 0.074 | 6.148 | 1.384 | < .001 |
| | Intermed., 90° | 0.338 | 0.060 | 5.641 | 1.025 | < .001 |
| | S/(L+M), 90° | 0.454 | 0.074 | 6.124 | 1.379 | < .001 |
| S/(L+M), 0° | L/(L+M), 18° | 0.181 | 0.074 | 2.447 | 0.551 | 1.000 |
| | Intermed., 18° | 0.101 | 0.074 | 1.355 | 0.305 | 1.000 |
| | S/(L+M), 18° | -0.009 | 0.060 | -0.155 | -0.028 | 1.000 |
| | L/(L+M), 36° | 0.218 | 0.074 | 2.943 | 0.663 | 0.646 |
| | Intermed., 36° | 0.124 | 0.074 | 1.668 | 0.376 | 1.000 |
| | S/(L+M), 36° | 0.060 | 0.060 | 0.996 | 0.181 | 1.000 |

| | | | | | | |
|---|---|---|---|---|---|---|
| | L/(L+M), 54° | 0.273 | 0.074 | 3.674 | 0.827 | 0.065 |
| | Intermed., 54° | 0.197 | 0.074 | 2.660 | 0.599 | 1.000 |
| | S/(L+M), 54° | 0.188 | 0.060 | 3.140 | 0.571 | 0.343 |
| | L/(L+M), 72° | 0.398 | 0.074 | 5.364 | 1.208 | < .001 |
| | Intermed., 72° | 0.311 | 0.074 | 4.189 | 0.943 | 0.011 |
| | S/(L+M), 72° | 0.363 | 0.060 | 6.063 | 1.102 | < .001 |
| | L/(L+M), 90° | 0.527 | 0.074 | 7.110 | 1.601 | < .001 |
| | Intermed., 90° | 0.409 | 0.074 | 5.515 | 1.242 | < .001 |
| | S/(L+M), 90° | 0.526 | 0.060 | 8.778 | 1.595 | < .001 |
| L/(L+M), 18° | Intermed., 18° | -0.081 | 0.060 | -1.346 | -0.246 | 1.000 |
| | S/(L+M), 18° | -0.191 | 0.060 | -3.172 | -0.579 | 0.396 |
| | L/(L+M), 36° | 0.037 | 0.060 | 0.615 | 0.112 | 1.000 |
| | Intermed., 36° | -0.058 | 0.074 | -0.779 | -0.175 | 1.000 |
| | S/(L+M), 36° | -0.122 | 0.074 | -1.643 | -0.370 | 1.000 |
| | L/(L+M), 54° | 0.091 | 0.060 | 1.521 | 0.276 | 1.000 |
| | Intermed., 54° | 0.016 | 0.074 | 0.213 | 0.048 | 1.000 |
| | S/(L+M), 54° | 0.007 | 0.074 | 0.088 | 0.020 | 1.000 |
| | L/(L+M), 72° | 0.216 | 0.060 | 3.614 | 0.657 | 0.074 |
| | Intermed., 72° | 0.129 | 0.074 | 1.742 | 0.392 | 1.000 |
| | S/(L+M), 72° | 0.181 | 0.074 | 2.447 | 0.551 | 1.000 |
| | L/(L+M), 90° | 0.346 | 0.060 | 5.777 | 1.050 | < .001 |
| | Intermed., 90° | 0.228 | 0.074 | 3.068 | 0.691 | 0.447 |
| | S/(L+M), 90° | 0.344 | 0.074 | 4.639 | 1.044 | 0.002 |
| Intermed., 18° | S/(L+M), 18° | -0.110 | 0.060 | -1.826 | -0.333 | 1.000 |
| | L/(L+M), 36° | 0.118 | 0.074 | 1.588 | 0.357 | 1.000 |
| | Intermed., 36° | 0.023 | 0.060 | 0.387 | 0.070 | 1.000 |
| | S/(L+M), 36° | -0.041 | 0.074 | -0.551 | -0.124 | 1.000 |
| | L/(L+M), 54° | 0.172 | 0.074 | 2.319 | 0.522 | 1.000 |
| | Intermed., 54° | 0.097 | 0.060 | 1.616 | 0.294 | 1.000 |
| | S/(L+M), 54° | 0.087 | 0.074 | 1.179 | 0.265 | 1.000 |
| | L/(L+M), 72° | 0.297 | 0.074 | 4.009 | 0.902 | 0.020 |
| | Intermed., 72° | 0.210 | 0.060 | 3.510 | 0.638 | 0.104 |
| | S/(L+M), 72° | 0.262 | 0.074 | 3.538 | 0.797 | 0.102 |
| | L/(L+M), 90° | 0.427 | 0.074 | 5.755 | 1.296 | < .001 |
| | Intermed., 90° | 0.309 | 0.060 | 5.153 | 0.936 | < .001 |
| | S/(L+M), 90° | 0.425 | 0.074 | 5.730 | 1.290 | < .001 |
| S/(L+M), 18° | L/(L+M), 36° | 0.228 | 0.074 | 3.069 | 0.691 | 0.446 |
| | Intermed., 36° | 0.133 | 0.074 | 1.793 | 0.404 | 1.000 |
| | S/(L+M), 36° | 0.069 | 0.060 | 1.152 | 0.209 | 1.000 |
| | L/(L+M), 54° | 0.282 | 0.074 | 3.800 | 0.855 | 0.042 |
| | Intermed., 54° | 0.207 | 0.074 | 2.785 | 0.627 | 1.000 |
| | S/(L+M), 54° | 0.197 | 0.060 | 3.295 | 0.599 | 0.210 |
| | L/(L+M), 72° | 0.407 | 0.074 | 5.489 | 1.236 | < .001 |
| | Intermed., 72° | 0.320 | 0.074 | 4.314 | 0.971 | 0.007 |
| | S/(L+M), 72° | 0.372 | 0.060 | 6.218 | 1.130 | < .001 |
| | L/(L+M), 90° | 0.537 | 0.074 | 7.235 | 1.629 | < .001 |

| | | | | | | |
|---|---|---|---|---|---|---|
| | Intermed., 90° | 0.418 | 0.074 | 5.640 | 1.270 | < .001 |
| | S/(L+M), 90° | 0.535 | 0.060 | 8.934 | 1.623 | < .001 |
| L/(L+M), 36° | Intermed., 36° | -0.095 | 0.060 | -1.573 | -0.287 | 1.000 |
| | S/(L+M), 36° | -0.159 | 0.060 | -2.638 | -0.482 | 1.000 |
| | L/(L+M), 54° | 0.054 | 0.060 | 0.906 | 0.165 | 1.000 |
| | Intermed., 54° | -0.021 | 0.074 | -0.283 | -0.064 | 1.000 |
| | S/(L+M), 54° | -0.030 | 0.074 | -0.409 | -0.092 | 1.000 |
| | L/(L+M), 72° | 0.180 | 0.060 | 2.999 | 0.545 | 0.527 |
| | Intermed., 72° | 0.092 | 0.074 | 1.245 | 0.280 | 1.000 |
| | S/(L+M), 72° | 0.145 | 0.074 | 1.950 | 0.439 | 1.000 |
| | L/(L+M), 90° | 0.309 | 0.060 | 5.162 | 0.938 | < .001 |
| | Intermed., 90° | 0.191 | 0.074 | 2.572 | 0.579 | 1.000 |
| | S/(L+M), 90° | 0.307 | 0.074 | 4.142 | 0.933 | 0.013 |
| Intermed., 36° | S/(L+M), 36° | -0.064 | 0.060 | -1.065 | -0.194 | 1.000 |
| | L/(L+M), 54° | 0.149 | 0.074 | 2.006 | 0.452 | 1.000 |
| | Intermed., 54° | 0.074 | 0.060 | 1.229 | 0.223 | 1.000 |
| | S/(L+M), 54° | 0.064 | 0.074 | 0.867 | 0.195 | 1.000 |
| | L/(L+M), 72° | 0.274 | 0.074 | 3.696 | 0.832 | 0.060 |
| | Intermed., 72° | 0.187 | 0.060 | 3.123 | 0.567 | 0.362 |
| | S/(L+M), 72° | 0.239 | 0.074 | 3.226 | 0.726 | 0.276 |
| | L/(L+M), 90° | 0.404 | 0.074 | 5.442 | 1.225 | < .001 |
| | Intermed., 90° | 0.285 | 0.060 | 4.766 | 0.866 | 0.001 |
| | S/(L+M), 90° | 0.402 | 0.074 | 5.418 | 1.220 | < .001 |
| S/(L+M), 36° | L/(L+M), 54° | 0.213 | 0.074 | 2.870 | 0.646 | 0.799 |
| | Intermed., 54° | 0.138 | 0.074 | 1.856 | 0.418 | 1.000 |
| | S/(L+M), 54° | 0.128 | 0.060 | 2.144 | 0.390 | 1.000 |
| | L/(L+M), 72° | 0.338 | 0.074 | 4.560 | 1.027 | 0.003 |
| | Intermed., 72° | 0.251 | 0.074 | 3.384 | 0.762 | 0.168 |
| | S/(L+M), 72° | 0.303 | 0.060 | 5.066 | 0.921 | < .001 |
| | L/(L+M), 90° | 0.468 | 0.074 | 6.306 | 1.420 | < .001 |
| | Intermed., 90° | 0.349 | 0.074 | 4.711 | 1.060 | 0.002 |
| | S/(L+M), 90° | 0.466 | 0.060 | 7.782 | 1.414 | < .001 |
| L/(L+M), 54° | Intermed., 54° | -0.075 | 0.060 | -1.251 | -0.228 | 1.000 |
| | S/(L+M), 54° | -0.085 | 0.060 | -1.406 | -0.257 | 1.000 |
| | L/(L+M), 72° | 0.125 | 0.060 | 2.093 | 0.380 | 1.000 |
| | Intermed., 72° | 0.038 | 0.074 | 0.514 | 0.116 | 1.000 |
| | S/(L+M), 72° | 0.090 | 0.074 | 1.219 | 0.275 | 1.000 |
| | L/(L+M), 90° | 0.255 | 0.060 | 4.256 | 0.773 | 0.007 |
| | Intermed., 90° | 0.137 | 0.074 | 1.841 | 0.414 | 1.000 |
| | S/(L+M), 90° | 0.253 | 0.074 | 3.411 | 0.768 | 0.154 |
| Intermed., 54° | S/(L+M), 54° | -0.009 | 0.060 | -0.155 | -0.028 | 1.000 |
| | L/(L+M), 72° | 0.201 | 0.074 | 2.704 | 0.609 | 1.000 |
| | Intermed., 72° | 0.113 | 0.060 | 1.894 | 0.344 | 1.000 |
| | S/(L+M), 72° | 0.166 | 0.074 | 2.234 | 0.503 | 1.000 |
| | L/(L+M), 90° | 0.330 | 0.074 | 4.450 | 1.002 | 0.004 |
| | Intermed., 90° | 0.212 | 0.060 | 3.537 | 0.643 | 0.096 |

| | | | | | | |
|---|---|---|---|---|---|---|
| | S/(L+M), 90° | 0.328 | 0.074 | 4.426 | 0.996 | 0.004 |
| S/(L+M), 54° | L/(L+M), 72° | 0.210 | 0.074 | 2.829 | 0.637 | 0.897 |
| | Intermed., 72° | 0.123 | 0.074 | 1.654 | 0.372 | 1.000 |
| | S/(L+M), 72° | 0.175 | 0.060 | 2.923 | 0.531 | 0.661 |
| | L/(L+M), 90° | 0.339 | 0.074 | 4.575 | 1.030 | 0.003 |
| | Intermed., 90° | 0.221 | 0.074 | 2.980 | 0.671 | 0.580 |
| | S/(L+M), 90° | 0.338 | 0.060 | 5.638 | 1.025 | < .001 |
| L/(L+M), 72° | Intermed., 72° | -0.087 | 0.060 | -1.450 | -0.265 | 1.000 |
| | S/(L+M), 72° | -0.035 | 0.060 | -0.580 | -0.106 | 1.000 |
| | L/(L+M), 90° | 0.129 | 0.060 | 2.163 | 0.393 | 1.000 |
| | Intermed., 90° | 0.011 | 0.074 | 0.151 | 0.034 | 1.000 |
| | S/(L+M), 90° | 0.128 | 0.074 | 1.722 | 0.388 | 1.000 |
| Intermed., 72° | S/(L+M), 72° | 0.052 | 0.060 | 0.870 | 0.159 | 1.000 |
| | L/(L+M), 90° | 0.217 | 0.074 | 2.921 | 0.658 | 0.689 |
| | Intermed., 90° | 0.098 | 0.060 | 1.643 | 0.299 | 1.000 |
| | S/(L+M), 90° | 0.215 | 0.074 | 2.897 | 0.652 | 0.739 |
| S/(L+M), 72° | L/(L+M), 90° | 0.164 | 0.074 | 2.216 | 0.499 | 1.000 |
| | Intermed., 90° | 0.046 | 0.074 | 0.621 | 0.140 | 1.000 |
| | S/(L+M), 90° | 0.163 | 0.060 | 2.716 | 0.493 | 1.000 |
| L/(L+M), 90° | Intermed., 90° | -0.118 | 0.060 | -1.967 | -0.359 | 1.000 |
| | S/(L+M), 90° | -0.002 | 0.060 | -0.030 | -0.005 | 1.000 |
| Intermed., 90° | S/(L+M), 90° | 0.117 | 0.060 | 1.937 | 0.354 | 1.000 |

*Note.* P-value adjusted for comparing a family of 153 estimates.

**Supplementary Table 2**

*Post hoc tests of interaction between axis and hue angle for $F_B$ (8 Hz) in Experiment 1*

| | | Mean Difference | SE | t | Cohen's d | $p_{bonf}$ |
|---|---|---|---|---|---|---|
| L/(L+M), 0° | Intermed., 0° | -0.156 | 0.067 | -2.315 | -0.648 | 1.000 |
| | S/(L+M), 0° | -0.247 | 0.067 | -3.657 | -1.023 | 0.125 |
| | L/(L+M), 18° | -0.020 | 0.042 | -0.471 | -0.082 | 1.000 |
| | Intermed., 18° | -0.114 | 0.070 | -1.629 | -0.473 | 1.000 |
| | S/(L+M), 18° | -0.220 | 0.070 | -3.139 | -0.911 | 0.481 |
| | L/(L+M), 36° | -0.038 | 0.042 | -0.895 | -0.156 | 1.000 |
| | Intermed., 36° | -0.115 | 0.070 | -1.639 | -0.476 | 1.000 |
| | S/(L+M), 36° | -0.295 | 0.070 | -4.208 | -1.222 | 0.021 |
| | L/(L+M), 54° | -0.098 | 0.042 | -2.325 | -0.405 | 1.000 |
| | Intermed., 54° | -0.141 | 0.070 | -2.011 | -0.584 | 1.000 |
| | S/(L+M), 54° | -0.336 | 0.070 | -4.798 | -1.393 | 0.003 |
| | L/(L+M), 72° | -0.124 | 0.042 | -2.941 | -0.512 | 0.587 |
| | Intermed., 72° | -0.220 | 0.070 | -3.141 | -0.912 | 0.479 |
| | S/(L+M), 72° | -0.394 | 0.070 | -5.625 | -1.633 | < .001 |
| | L/(L+M), 90° | -0.212 | 0.042 | -5.054 | -0.880 | < .001 |
| | Intermed., 90° | -0.322 | 0.070 | -4.597 | -1.334 | 0.006 |
| | S/(L+M), 90° | -0.602 | 0.070 | -8.587 | -2.493 | < .001 |
| Intermed., 0° | S/(L+M), 0° | -0.091 | 0.067 | -1.342 | -0.375 | 1.000 |

| | | | | | | |
|---|---|---|---|---|---|---|
| | L/(L+M), 18° | 0.136 | 0.070 | 1.948 | 0.566 | 1.000 |
| | Intermed., 18° | 0.042 | 0.042 | 1.003 | 0.175 | 1.000 |
| | S/(L+M), 18° | -0.064 | 0.070 | -0.908 | -0.264 | 1.000 |
| | L/(L+M), 36° | 0.119 | 0.070 | 1.694 | 0.492 | 1.000 |
| | Intermed., 36° | 0.041 | 0.042 | 0.987 | 0.172 | 1.000 |
| | S/(L+M), 36° | -0.139 | 0.070 | -1.978 | -0.574 | 1.000 |
| | L/(L+M), 54° | 0.059 | 0.070 | 0.836 | 0.243 | 1.000 |
| | Intermed., 54° | 0.015 | 0.042 | 0.366 | 0.064 | 1.000 |
| | S/(L+M), 54° | -0.180 | 0.070 | -2.567 | -0.745 | 1.000 |
| | L/(L+M), 72° | 0.033 | 0.070 | 0.467 | 0.136 | 1.000 |
| | Intermed., 72° | -0.064 | 0.042 | -1.517 | -0.264 | 1.000 |
| | S/(L+M), 72° | -0.238 | 0.070 | -3.395 | -0.985 | 0.235 |
| | L/(L+M), 90° | -0.056 | 0.070 | -0.801 | -0.232 | 1.000 |
| | Intermed., 90° | -0.166 | 0.042 | -3.944 | -0.687 | 0.019 |
| | S/(L+M), 90° | -0.445 | 0.070 | -6.356 | -1.845 | < .001 |
| S/(L+M), 0° | L/(L+M), 18° | 0.227 | 0.070 | 3.241 | 0.941 | 0.363 |
| | Intermed., 18° | 0.133 | 0.070 | 1.894 | 0.550 | 1.000 |
| | S/(L+M), 18° | 0.027 | 0.042 | 0.641 | 0.112 | 1.000 |
| | L/(L+M), 36° | 0.209 | 0.070 | 2.987 | 0.867 | 0.727 |
| | Intermed., 36° | 0.132 | 0.070 | 1.885 | 0.547 | 1.000 |
| | S/(L+M), 36° | -0.048 | 0.042 | -1.142 | -0.199 | 1.000 |
| | L/(L+M), 54° | 0.149 | 0.070 | 2.129 | 0.618 | 1.000 |
| | Intermed., 54° | 0.106 | 0.070 | 1.513 | 0.439 | 1.000 |
| | S/(L+M), 54° | -0.089 | 0.042 | -2.125 | -0.370 | 1.000 |
| | L/(L+M), 72° | 0.123 | 0.070 | 1.760 | 0.511 | 1.000 |
| | Intermed., 72° | 0.027 | 0.070 | 0.383 | 0.111 | 1.000 |
| | S/(L+M), 72° | -0.147 | 0.042 | -3.504 | -0.610 | 0.094 |
| | L/(L+M), 90° | 0.034 | 0.070 | 0.492 | 0.143 | 1.000 |
| | Intermed., 90° | -0.075 | 0.070 | -1.073 | -0.311 | 1.000 |
| | S/(L+M), 90° | -0.355 | 0.042 | -8.442 | -1.470 | < .001 |
| L/(L+M), 18° | Intermed., 18° | -0.094 | 0.067 | -1.398 | -0.391 | 1.000 |
| | S/(L+M), 18° | -0.200 | 0.067 | -2.964 | -0.829 | 0.824 |
| | L/(L+M), 36° | -0.018 | 0.042 | -0.424 | -0.074 | 1.000 |
| | Intermed., 36° | -0.095 | 0.070 | -1.357 | -0.394 | 1.000 |
| | S/(L+M), 36° | -0.275 | 0.070 | -3.926 | -1.140 | 0.050 |
| | L/(L+M), 54° | -0.078 | 0.042 | -1.854 | -0.323 | 1.000 |
| | Intermed., 54° | -0.121 | 0.070 | -1.729 | -0.502 | 1.000 |
| | S/(L+M), 54° | -0.316 | 0.070 | -4.516 | -1.311 | 0.008 |
| | L/(L+M), 72° | -0.104 | 0.042 | -2.470 | -0.430 | 1.000 |
| | Intermed., 72° | -0.200 | 0.070 | -2.858 | -0.830 | 1.000 |
| | S/(L+M), 72° | -0.374 | 0.070 | -5.343 | -1.551 | < .001 |
| | L/(L+M), 90° | -0.193 | 0.042 | -4.583 | -0.798 | 0.002 |
| | Intermed., 90° | -0.302 | 0.070 | -4.314 | -1.252 | 0.015 |
| | S/(L+M), 90° | -0.582 | 0.070 | -8.305 | -2.411 | < .001 |
| Intermed., 18° | S/(L+M), 18° | -0.106 | 0.067 | -1.567 | -0.438 | 1.000 |
| | L/(L+M), 36° | 0.077 | 0.070 | 1.092 | 0.317 | 1.000 |

| | | | | | | |
|---|---|---|---|---|---|---|
| | Intermed., 36° | >0.001 | 0.042 | -0.016 | -0.003 | 1.000 |
| | S/(L+M), 36° | -0.181 | 0.070 | -2.579 | -0.749 | 1.000 |
| | L/(L+M), 54° | 0.016 | 0.070 | 0.234 | 0.068 | 1.000 |
| | Intermed., 54° | -0.027 | 0.042 | -0.637 | -0.111 | 1.000 |
| | S/(L+M), 54° | -0.222 | 0.070 | -3.169 | -0.920 | 0.443 |
| | L/(L+M), 72° | -0.009 | 0.070 | -0.135 | -0.039 | 1.000 |
| | Intermed., 72° | -0.106 | 0.042 | -2.520 | -0.439 | 1.000 |
| | S/(L+M), 72° | -0.280 | 0.070 | -3.996 | -1.160 | 0.040 |
| | L/(L+M), 90° | -0.098 | 0.070 | -1.402 | -0.407 | 1.000 |
| | Intermed., 90° | -0.208 | 0.042 | -4.947 | -0.861 | < .001 |
| | S/(L+M), 90° | -0.487 | 0.070 | -6.958 | -2.020 | < .001 |
| S/(L+M), 18° | L/(L+M), 36° | 0.182 | 0.070 | 2.602 | 0.755 | 1.000 |
| | Intermed., 36° | 0.105 | 0.070 | 1.500 | 0.435 | 1.000 |
| | S/(L+M), 36° | -0.075 | 0.042 | -1.783 | -0.310 | 1.000 |
| | L/(L+M), 54° | 0.122 | 0.070 | 1.744 | 0.506 | 1.000 |
| | Intermed., 54° | 0.079 | 0.070 | 1.128 | 0.327 | 1.000 |
| | S/(L+M), 54° | -0.116 | 0.042 | -2.766 | -0.482 | 0.986 |
| | L/(L+M), 72° | 0.096 | 0.070 | 1.375 | 0.399 | 1.000 |
| | Intermed., 72° | < -0.001 | 0.070 | -0.002 | < -0.001 | 1.000 |
| | S/(L+M), 72° | -0.174 | 0.042 | -4.146 | -0.722 | 0.009 |
| | L/(L+M), 90° | 0.008 | 0.070 | 0.107 | 0.031 | 1.000 |
| | Intermed., 90° | -0.102 | 0.070 | -1.458 | -0.423 | 1.000 |
| | S/(L+M), 90° | -0.382 | 0.042 | -9.084 | -1.582 | < .001 |
| L/(L+M), 36° | Intermed., 36° | -0.077 | 0.067 | -1.144 | -0.320 | 1.000 |
| | S/(L+M), 36° | -0.257 | 0.067 | -3.810 | -1.066 | 0.081 |
| | L/(L+M), 54° | -0.060 | 0.042 | -1.430 | -0.249 | 1.000 |
| | Intermed., 54° | -0.103 | 0.070 | -1.474 | -0.428 | 1.000 |
| | S/(L+M), 54° | -0.298 | 0.070 | -4.261 | -1.237 | 0.018 |
| | L/(L+M), 72° | -0.086 | 0.042 | -2.045 | -0.356 | 1.000 |
| | Intermed., 72° | -0.182 | 0.070 | -2.603 | -0.756 | 1.000 |
| | S/(L+M), 72° | -0.356 | 0.070 | -5.088 | -1.477 | 0.001 |
| | L/(L+M), 90° | -0.175 | 0.042 | -4.159 | -0.724 | 0.009 |
| | Intermed., 90° | -0.284 | 0.070 | -4.059 | -1.178 | 0.033 |
| | S/(L+M), 90° | -0.564 | 0.070 | -8.050 | -2.337 | < .001 |
| Intermed., 36° | S/(L+M), 36° | -0.180 | 0.067 | -2.667 | -0.746 | 1.000 |
| | L/(L+M), 54° | 0.017 | 0.070 | 0.244 | 0.071 | 1.000 |
| | Intermed., 54° | -0.026 | 0.042 | -0.620 | -0.108 | 1.000 |
| | S/(L+M), 54° | -0.221 | 0.070 | -3.159 | -0.917 | 0.455 |
| | L/(L+M), 72° | -0.009 | 0.070 | -0.125 | -0.036 | 1.000 |
| | Intermed., 72° | -0.105 | 0.042 | -2.503 | -0.436 | 1.000 |
| | S/(L+M), 72° | -0.279 | 0.070 | -3.986 | -1.157 | 0.041 |
| | L/(L+M), 90° | -0.098 | 0.070 | -1.392 | -0.404 | 1.000 |
| | Intermed., 90° | -0.207 | 0.042 | -4.931 | -0.859 | < .001 |
| | S/(L+M), 90° | -0.487 | 0.070 | -6.948 | -2.017 | < .001 |
| S/(L+M), 36° | L/(L+M), 54° | 0.197 | 0.070 | 2.814 | 0.817 | 1.000 |
| | Intermed., 54° | 0.154 | 0.070 | 2.197 | 0.638 | 1.000 |

| | | | | | | |
|---|---|---|---|---|---|---|
| | S/(L+M), 54° | -0.041 | 0.042 | -0.983 | -0.171 | 1.000 |
| | L/(L+M), 72° | 0.171 | 0.070 | 2.444 | 0.710 | 1.000 |
| | Intermed., 72° | 0.075 | 0.070 | 1.068 | 0.310 | 1.000 |
| | S/(L+M), 72° | -0.099 | 0.042 | -2.363 | -0.411 | 1.000 |
| | L/(L+M), 90° | 0.082 | 0.070 | 1.177 | 0.342 | 1.000 |
| | Intermed., 90° | -0.027 | 0.070 | -0.388 | -0.113 | 1.000 |
| | S/(L+M), 90° | -0.307 | 0.042 | -7.300 | -1.271 | < .001 |
| L/(L+M), 54° | Intermed., 54° | -0.043 | 0.067 | -0.640 | -0.179 | 1.000 |
| | S/(L+M), 54° | -0.238 | 0.067 | -3.532 | -0.988 | 0.178 |
| | L/(L+M), 72° | -0.026 | 0.042 | -0.615 | -0.107 | 1.000 |
| | Intermed., 72° | -0.122 | 0.070 | -1.746 | -0.507 | 1.000 |
| | S/(L+M), 72° | -0.296 | 0.070 | -4.231 | -1.228 | 0.020 |
| | L/(L+M), 90° | -0.115 | 0.042 | -2.728 | -0.475 | 1.000 |
| | Intermed., 90° | -0.224 | 0.070 | -3.202 | -0.929 | 0.404 |
| | S/(L+M), 90° | -0.504 | 0.070 | -7.192 | -2.088 | < .001 |
| Intermed., 54° | S/(L+M), 54° | -0.195 | 0.067 | -2.892 | -0.809 | 0.993 |
| | L/(L+M), 72° | 0.017 | 0.070 | 0.247 | 0.072 | 1.000 |
| | Intermed., 72° | -0.079 | 0.042 | -1.883 | -0.328 | 1.000 |
| | S/(L+M), 72° | -0.253 | 0.070 | -3.614 | -1.049 | 0.125 |
| | L/(L+M), 90° | -0.071 | 0.070 | -1.020 | -0.296 | 1.000 |
| | Intermed., 90° | -0.181 | 0.042 | -4.311 | -0.751 | 0.005 |
| | S/(L+M), 90° | -0.461 | 0.070 | -6.576 | -1.909 | < .001 |
| S/(L+M), 54° | L/(L+M), 72° | 0.213 | 0.070 | 3.034 | 0.881 | 0.640 |
| | Intermed., 72° | 0.116 | 0.070 | 1.657 | 0.481 | 1.000 |
| | S/(L+M), 72° | -0.058 | 0.042 | -1.379 | -0.240 | 1.000 |
| | L/(L+M), 90° | 0.124 | 0.070 | 1.767 | 0.513 | 1.000 |
| | Intermed., 90° | 0.014 | 0.070 | 0.201 | 0.058 | 1.000 |
| | S/(L+M), 90° | -0.265 | 0.042 | -6.317 | -1.100 | < .001 |
| L/(L+M), 72° | Intermed., 72° | -0.096 | 0.067 | -1.429 | -0.400 | 1.000 |
| | S/(L+M), 72° | -0.271 | 0.067 | -4.008 | -1.121 | 0.046 |
| | L/(L+M), 90° | -0.089 | 0.042 | -2.113 | -0.368 | 1.000 |
| | Intermed., 90° | -0.198 | 0.070 | -2.833 | -0.822 | 1.000 |
| | S/(L+M), 90° | -0.478 | 0.070 | -6.823 | -1.981 | < .001 |
| Intermed., 72° | S/(L+M), 72° | -0.174 | 0.067 | -2.579 | -0.721 | 1.000 |
| | L/(L+M), 90° | 0.008 | 0.070 | 0.109 | 0.032 | 1.000 |
| | Intermed., 90° | -0.102 | 0.042 | -2.427 | -0.423 | 1.000 |
| | S/(L+M), 90° | -0.382 | 0.070 | -5.447 | -1.581 | < .001 |
| S/(L+M), 72° | L/(L+M), 90° | 0.182 | 0.070 | 2.594 | 0.753 | 1.000 |
| | Intermed., 90° | 0.072 | 0.070 | 1.029 | 0.299 | 1.000 |
| | S/(L+M), 90° | -0.207 | 0.042 | -4.938 | -0.860 | < .001 |
| L/(L+M), 90° | Intermed., 90° | -0.110 | 0.067 | -1.624 | -0.454 | 1.000 |
| | S/(L+M), 90° | -0.389 | 0.067 | -5.766 | -1.613 | < .001 |
| Intermed., 90° | S/(L+M), 90° | -0.280 | 0.067 | -4.142 | -1.158 | 0.031 |

*Note.* P-value adjusted for comparing a family of 153 estimates.

Experiment 2

**Supplementary Table 3**

*Post hoc tests of hue angle for $I_1$ (14.66 Hz) in Experiment 2*

| | | Mean Difference | SE | t | Cohen's d | $p_{bonf}$ |
|---|---|---|---|---|---|---|
| 0° | 18° | 0.022 | 0.075 | 0.290 | 0.056 | 1.000 |
| | 36° | 0.063 | 0.075 | 0.834 | 0.160 | 1.000 |
| | 54° | 0.186 | 0.075 | 2.478 | 0.477 | 0.249 |
| | 72° | 0.308 | 0.075 | 4.098 | 0.788 | 0.002 |
| | 90° | 0.402 | 0.075 | 5.343 | 1.028 | < .001 |
| 18° | 36° | 0.041 | 0.075 | 0.544 | 0.105 | 1.000 |
| | 54° | 0.165 | 0.075 | 2.188 | 0.421 | 0.501 |
| | 72° | 0.286 | 0.075 | 3.808 | 0.733 | 0.006 |
| | 90° | 0.380 | 0.075 | 5.053 | 0.972 | < .001 |
| 36° | 54° | 0.124 | 0.075 | 1.644 | 0.316 | 1.000 |
| | 72° | 0.245 | 0.075 | 3.264 | 0.628 | 0.030 |
| | 90° | 0.339 | 0.075 | 4.509 | 0.867 | < .001 |
| 54° | 72° | 0.122 | 0.075 | 1.620 | 0.312 | 1.000 |
| | 90° | 0.215 | 0.075 | 2.865 | 0.551 | 0.091 |
| 72° | 90° | 0.094 | 0.075 | 1.245 | 0.240 | 1.000 |

*Note.* P-value adjusted for comparing a family of 15 estimates.

**Supplementary Table 4**

*Post hoc tests of hue angle for $F_B$ (8 Hz) in Experiment 2*

| | | Mean Difference | SE | t | Cohen's d | $p_{bonf}$ |
|---|---|---|---|---|---|---|
| 0° | 18° | -0.057 | 0.049 | -1.162 | -0.095 | 1.000 |
| | 36° | -0.055 | 0.049 | -1.125 | -0.092 | 1.000 |
| | 54° | -0.035 | 0.049 | -0.719 | -0.059 | 1.000 |
| | 72° | -0.009 | 0.049 | -0.184 | -0.015 | 1.000 |
| | 90° | -0.022 | 0.049 | -0.443 | -0.036 | 1.000 |
| 18° | 36° | 0.002 | 0.049 | 0.037 | 0.003 | 1.000 |
| | 54° | 0.022 | 0.049 | 0.443 | 0.036 | 1.000 |
| | 72° | 0.048 | 0.049 | 0.977 | 0.080 | 1.000 |
| | 90° | 0.035 | 0.049 | 0.719 | 0.059 | 1.000 |
| 36° | 54° | 0.020 | 0.049 | 0.406 | 0.033 | 1.000 |
| | 72° | 0.046 | 0.049 | 0.941 | 0.077 | 1.000 |
| | 90° | 0.034 | 0.049 | 0.682 | 0.056 | 1.000 |
| 54° | 72° | 0.026 | 0.049 | 0.535 | 0.044 | 1.000 |
| | 90° | 0.014 | 0.049 | 0.277 | 0.023 | 1.000 |
| 72° | 90° | -0.013 | 0.049 | -0.258 | -0.021 | 1.000 |

*Note.* P-value adjusted for comparing a family of 15 estimates.

Experiment 3

**Supplementary Table 5**

*Post hoc tests of interaction between size and hue angle for $I_1$ (14.66 Hz) in Experiment 3*

| | | Mean Difference | SE | t | Cohen's d | $p_{bonf}$ |
|---|---|---|---|---|---|---|
| .096°, 0° | .4°, 0° | 0.019 | 0.086 | 0.222 | 0.054 | 1.000 |
| | 1.3°, 0° | -0.212 | 0.086 | -2.462 | -0.597 | 1.000 |

| | | | | | | |
|---|---|---|---|---|---|---|
| | 4.8°, 0° | 0.011 | 0.086 | 0.127 | 0.031 | 1.000 |
| | .096°, 30° | 0.076 | 0.090 | 0.847 | 0.215 | 1.000 |
| | .4°, 30° | 0.135 | 0.102 | 1.325 | 0.382 | 1.000 |
| | 1.3°, 30° | -0.115 | 0.102 | -1.120 | -0.323 | 1.000 |
| | 4.8°, 30° | 0.082 | 0.102 | 0.800 | 0.231 | 1.000 |
| | .096°, 60° | 0.365 | 0.090 | 4.055 | 1.030 | 0.013 |
| | .4°, 60° | 0.396 | 0.102 | 3.876 | 1.117 | 0.023 |
| | 1.3°, 60° | 0.178 | 0.102 | 1.743 | 0.502 | 1.000 |
| | 4.8°, 60° | 0.306 | 0.102 | 2.996 | 0.863 | 0.416 |
| | .096°, 90° | 0.554 | 0.090 | 6.143 | 1.560 | < .001 |
| | .4°, 90° | 0.485 | 0.102 | 4.748 | 1.368 | < .001 |
| | 1.3°, 90° | 0.527 | 0.102 | 5.157 | 1.485 | < .001 |
| | 4.8°, 90° | 0.636 | 0.102 | 6.223 | 1.793 | < .001 |
| .4°, 0° | 1.3°, 0° | -0.231 | 0.086 | -2.684 | -0.651 | 1.000 |
| | 4.8°, 0° | -0.008 | 0.086 | -0.095 | -0.023 | 1.000 |
| | .096°, 30° | 0.057 | 0.102 | 0.560 | 0.161 | 1.000 |
| | .4°, 30° | 0.116 | 0.090 | 1.291 | 0.328 | 1.000 |
| | 1.3°, 30° | -0.134 | 0.102 | -1.307 | -0.376 | 1.000 |
| | 4.8°, 30° | 0.063 | 0.102 | 0.613 | 0.177 | 1.000 |
| | .096°, 60° | 0.346 | 0.102 | 3.387 | 0.976 | 0.122 |
| | .4°, 60° | 0.377 | 0.090 | 4.186 | 1.063 | 0.008 |
| | 1.3°, 60° | 0.159 | 0.102 | 1.556 | 0.448 | 1.000 |
| | 4.8°, 60° | 0.287 | 0.102 | 2.809 | 0.809 | 0.720 |
| | .096°, 90° | 0.535 | 0.102 | 5.228 | 1.506 | < .001 |
| | .4°, 90° | 0.466 | 0.090 | 5.174 | 1.314 | < .001 |
| | 1.3°, 90° | 0.508 | 0.102 | 4.970 | 1.432 | < .001 |
| | 4.8°, 90° | 0.617 | 0.102 | 6.037 | 1.739 | < .001 |
| 1.3°, 0° | 4.8°, 0° | 0.223 | 0.086 | 2.589 | 0.627 | 1.000 |
| | .096°, 30° | 0.288 | 0.102 | 2.818 | 0.812 | 0.702 |
| | .4°, 30° | 0.347 | 0.102 | 3.396 | 0.978 | 0.119 |
| | 1.3°, 30° | 0.097 | 0.090 | 1.079 | 0.274 | 1.000 |
| | 4.8°, 30° | 0.294 | 0.102 | 2.872 | 0.827 | 0.602 |
| | .096°, 60° | 0.577 | 0.102 | 5.646 | 1.626 | < .001 |
| | .4°, 60° | 0.608 | 0.102 | 5.948 | 1.713 | < .001 |
| | 1.3°, 60° | 0.390 | 0.090 | 4.327 | 1.099 | 0.005 |
| | 4.8°, 60° | 0.518 | 0.102 | 5.068 | 1.460 | < .001 |
| | .096°, 90° | 0.765 | 0.102 | 7.486 | 2.156 | < .001 |
| | .4°, 90° | 0.697 | 0.102 | 6.819 | 1.964 | < .001 |
| | 1.3°, 90° | 0.739 | 0.090 | 8.200 | 2.082 | < .001 |
| | 4.8°, 90° | 0.848 | 0.102 | 8.295 | 2.390 | < .001 |
| 4.8°, 0° | .096°, 30° | 0.065 | 0.102 | 0.640 | 0.184 | 1.000 |
| | .4°, 30° | 0.125 | 0.102 | 1.218 | 0.351 | 1.000 |
| | 1.3°, 30° | -0.125 | 0.102 | -1.227 | -0.353 | 1.000 |
| | 4.8°, 30° | 0.071 | 0.090 | 0.787 | 0.200 | 1.000 |
| | .096°, 60° | 0.355 | 0.102 | 3.467 | 0.999 | 0.094 |
| | .4°, 60° | 0.385 | 0.102 | 3.770 | 1.086 | 0.034 |

| | | | | | | |
|---|---|---|---|---|---|---|
| | 1.3°, 60° | 0.167 | 0.102 | 1.636 | 0.471 | 1.000 |
| | 4.8°, 60° | 0.295 | 0.090 | 3.278 | 0.832 | 0.182 |
| | .096°, 90° | 0.543 | 0.102 | 5.308 | 1.529 | < .001 |
| | .4°, 90° | 0.475 | 0.102 | 4.641 | 1.337 | 0.001 |
| | 1.3°, 90° | 0.516 | 0.102 | 5.050 | 1.455 | < .001 |
| | 4.8°, 90° | 0.625 | 0.090 | 6.940 | 1.762 | < .001 |
| .096°, 30° | .4°, 30° | 0.059 | 0.086 | 0.687 | 0.166 | 1.000 |
| | 1.3°, 30° | -0.191 | 0.086 | -2.219 | -0.538 | 1.000 |
| | 4.8°, 30° | 0.005 | 0.086 | 0.063 | 0.015 | 1.000 |
| | .096°, 60° | 0.289 | 0.090 | 3.208 | 0.814 | 0.227 |
| | .4°, 60° | 0.320 | 0.102 | 3.130 | 0.902 | 0.277 |
| | 1.3°, 60° | 0.102 | 0.102 | 0.996 | 0.287 | 1.000 |
| | 4.8°, 60° | 0.230 | 0.102 | 2.249 | 0.648 | 1.000 |
| | .096°, 90° | 0.477 | 0.090 | 5.295 | 1.345 | < .001 |
| | .4°, 90° | 0.409 | 0.102 | 4.001 | 1.153 | 0.015 |
| | 1.3°, 90° | 0.451 | 0.102 | 4.410 | 1.270 | 0.003 |
| | 4.8°, 90° | 0.560 | 0.102 | 5.477 | 1.578 | < .001 |
| .4°, 30° | 1.3°, 30° | -0.250 | 0.086 | -2.906 | -0.704 | 0.550 |
| | 4.8°, 30° | -0.054 | 0.086 | -0.623 | -0.151 | 1.000 |
| | .096°, 60° | 0.230 | 0.102 | 2.249 | 0.648 | 1.000 |
| | .4°, 60° | 0.261 | 0.090 | 2.895 | 0.735 | 0.580 |
| | 1.3°, 60° | 0.043 | 0.102 | 0.418 | 0.120 | 1.000 |
| | 4.8°, 60° | 0.171 | 0.102 | 1.671 | 0.481 | 1.000 |
| | .096°, 90° | 0.418 | 0.102 | 4.090 | 1.178 | 0.011 |
| | .4°, 90° | 0.350 | 0.090 | 3.883 | 0.986 | 0.025 |
| | 1.3°, 90° | 0.392 | 0.102 | 3.832 | 1.104 | 0.027 |
| | 4.8°, 90° | 0.501 | 0.102 | 4.899 | 1.411 | < .001 |
| 1.3°, 30° | 4.8°, 30° | 0.196 | 0.086 | 2.282 | 0.553 | 1.000 |
| | .096°, 60° | 0.480 | 0.102 | 4.694 | 1.352 | 0.001 |
| | .4°, 60° | 0.511 | 0.102 | 4.997 | 1.439 | < .001 |
| | 1.3°, 60° | 0.293 | 0.090 | 3.248 | 0.825 | 0.200 |
| | 4.8°, 60° | 0.421 | 0.102 | 4.116 | 1.186 | 0.010 |
| | .096°, 90° | 0.668 | 0.102 | 6.535 | 1.882 | < .001 |
| | .4°, 90° | 0.600 | 0.102 | 5.868 | 1.690 | < .001 |
| | 1.3°, 90° | 0.642 | 0.090 | 7.121 | 1.808 | < .001 |
| | 4.8°, 90° | 0.751 | 0.102 | 7.344 | 2.115 | < .001 |
| 4.8°, 30° | .096°, 60° | 0.284 | 0.102 | 2.774 | 0.799 | 0.797 |
| | .4°, 60° | 0.315 | 0.102 | 3.076 | 0.886 | 0.327 |
| | 1.3°, 60° | 0.096 | 0.102 | 0.942 | 0.271 | 1.000 |
| | 4.8°, 60° | 0.225 | 0.090 | 2.491 | 0.633 | 1.000 |
| | .096°, 90° | 0.472 | 0.102 | 4.614 | 1.329 | 0.001 |
| | .4°, 90° | 0.404 | 0.102 | 3.947 | 1.137 | 0.018 |
| | 1.3°, 90° | 0.445 | 0.102 | 4.356 | 1.255 | 0.004 |
| | 4.8°, 90° | 0.555 | 0.090 | 6.153 | 1.562 | < .001 |
| .096°, 60° | .4°, 60° | 0.031 | 0.086 | 0.359 | 0.087 | 1.000 |
| | 1.3°, 60° | -0.187 | 0.086 | -2.177 | -0.528 | 1.000 |

| | | | | | | |
|---|---|---|---|---|---|---|
| | 4.8°, 60° | -0.059 | 0.086 | -0.687 | -0.166 | 1.000 |
| | .096°, 90° | 0.188 | 0.090 | 2.088 | 0.530 | 1.000 |
| | .4°, 90° | 0.120 | 0.102 | 1.174 | 0.338 | 1.000 |
| | 1.3°, 90° | 0.162 | 0.102 | 1.583 | 0.456 | 1.000 |
| | 4.8°, 90° | 0.271 | 0.102 | 2.649 | 0.763 | 1.000 |
| .4°, 60° | 1.3°, 60° | -0.218 | 0.086 | -2.536 | -0.615 | 1.000 |
| | 4.8°, 60° | -0.090 | 0.086 | -1.046 | -0.254 | 1.000 |
| | .096°, 90° | 0.157 | 0.102 | 1.538 | 0.443 | 1.000 |
| | .4°, 90° | 0.089 | 0.090 | 0.988 | 0.251 | 1.000 |
| | 1.3°, 90° | 0.131 | 0.102 | 1.280 | 0.369 | 1.000 |
| | 4.8°, 90° | 0.240 | 0.102 | 2.347 | 0.676 | 1.000 |
| 1.3°, 60° | 4.8°, 60° | 0.128 | 0.086 | 1.490 | 0.361 | 1.000 |
| | .096°, 90° | 0.375 | 0.102 | 3.672 | 1.058 | 0.047 |
| | .4°, 90° | 0.307 | 0.102 | 3.005 | 0.866 | 0.405 |
| | 1.3°, 90° | 0.349 | 0.090 | 3.873 | 0.983 | 0.025 |
| | 4.8°, 90° | 0.458 | 0.102 | 4.481 | 1.291 | 0.002 |
| 4.8°, 60° | .096°, 90° | 0.247 | 0.102 | 2.418 | 0.697 | 1.000 |
| | .4°, 90° | 0.179 | 0.102 | 1.751 | 0.505 | 1.000 |
| | 1.3°, 90° | 0.221 | 0.102 | 2.160 | 0.622 | 1.000 |
| | 4.8°, 90° | 0.330 | 0.090 | 3.661 | 0.930 | 0.052 |
| .096°, 90° | .4°, 90° | -0.068 | 0.086 | -0.793 | -0.192 | 1.000 |
| | 1.3°, 90° | -0.026 | 0.086 | -0.306 | -0.074 | 1.000 |
| | 4.8°, 90° | 0.083 | 0.086 | 0.962 | 0.233 | 1.000 |
| .4°, 90° | 1.3°, 90° | 0.042 | 0.086 | 0.486 | 0.118 | 1.000 |
| | 4.8°, 90° | 0.151 | 0.086 | 1.754 | 0.425 | 1.000 |
| 1.3°, 90° | 4.8°, 90° | 0.109 | 0.086 | 1.268 | 0.307 | 1.000 |

*Note.* P-value adjusted for comparing a family of 120 estimates.

**Supplementary Table 6**

*Post hoc tests of interaction between size and hue angle for $F_B$ (8 Hz) in Experiment 3*

| | | Mean Difference | SE | t | Cohen's d | $p_{bonf}$ |
|---|---|---|---|---|---|---|
| .096°, 0° | .4°, 0° | 0.263 | 0.152 | 1.726 | 0.490 | 1.000 |
| | 1.3°, 0° | 0.214 | 0.152 | 1.403 | 0.398 | 1.000 |
| | 4.8°, 0° | 0.090 | 0.152 | 0.591 | 0.168 | 1.000 |
| | .096°, 30° | 0.074 | 0.159 | 0.463 | 0.137 | 1.000 |
| | .4°, 30° | 0.085 | 0.164 | 0.522 | 0.159 | 1.000 |
| | 1.3°, 30° | 0.320 | 0.164 | 1.956 | 0.596 | 1.000 |
| | 4.8°, 30° | 0.155 | 0.164 | 0.950 | 0.290 | 1.000 |
| | .096°, 60° | -0.065 | 0.159 | -0.411 | -0.122 | 1.000 |
| | .4°, 60° | -0.065 | 0.164 | -0.400 | -0.122 | 1.000 |
| | 1.3°, 60° | -0.025 | 0.164 | -0.150 | -0.046 | 1.000 |
| | 4.8°, 60° | 0.132 | 0.164 | 0.806 | 0.246 | 1.000 |
| | .096°, 90° | 0.003 | 0.159 | 0.017 | 0.005 | 1.000 |
| | .4°, 90° | -0.452 | 0.164 | -2.762 | -0.842 | 0.792 |
| | 1.3°, 90° | -0.651 | 0.164 | -3.978 | -1.213 | 0.014 |

| | | | | | | |
|---|---|---|---|---|---|---|
| | 4.8°, 90° | -0.215 | 0.164 | -1.311 | -0.400 | 1.000 |
| .4°, 0° | 1.3°, 0° | -0.049 | 0.152 | -0.322 | -0.091 | 1.000 |
| | 4.8°, 0° | -0.173 | 0.152 | -1.135 | -0.322 | 1.000 |
| | .096°, 30° | -0.189 | 0.164 | -1.156 | -0.352 | 1.000 |
| | .4°, 30° | -0.177 | 0.159 | -1.114 | -0.330 | 1.000 |
| | 1.3°, 30° | 0.057 | 0.164 | 0.350 | 0.107 | 1.000 |
| | 4.8°, 30° | -0.107 | 0.164 | -0.656 | -0.200 | 1.000 |
| | .096°, 60° | -0.328 | 0.164 | -2.006 | -0.612 | 1.000 |
| | .4°, 60° | -0.328 | 0.159 | -2.062 | -0.612 | 1.000 |
| | 1.3°, 60° | -0.287 | 0.164 | -1.756 | -0.535 | 1.000 |
| | 4.8°, 60° | -0.131 | 0.164 | -0.800 | -0.244 | 1.000 |
| | .096°, 90° | -0.260 | 0.164 | -1.589 | -0.485 | 1.000 |
| | .4°, 90° | -0.715 | 0.159 | -4.490 | -1.332 | 0.002 |
| | 1.3°, 90° | -0.914 | 0.164 | -5.584 | -1.703 | < .001 |
| | 4.8°, 90° | -0.477 | 0.164 | -2.917 | -0.889 | 0.501 |
| 1.3°, 0° | 4.8°, 0° | -0.124 | 0.152 | -0.812 | -0.230 | 1.000 |
| | .096°, 30° | -0.140 | 0.164 | -0.856 | -0.261 | 1.000 |
| | .4°, 30° | -0.128 | 0.164 | -0.783 | -0.239 | 1.000 |
| | 1.3°, 30° | 0.106 | 0.159 | 0.668 | 0.198 | 1.000 |
| | 4.8°, 30° | -0.058 | 0.164 | -0.356 | -0.108 | 1.000 |
| | .096°, 60° | -0.279 | 0.164 | -1.706 | -0.520 | 1.000 |
| | .4°, 60° | -0.279 | 0.164 | -1.706 | -0.520 | 1.000 |
| | 1.3°, 60° | -0.238 | 0.159 | -1.497 | -0.444 | 1.000 |
| | 4.8°, 60° | -0.082 | 0.164 | -0.500 | -0.152 | 1.000 |
| | .096°, 90° | -0.211 | 0.164 | -1.289 | -0.393 | 1.000 |
| | .4°, 90° | -0.665 | 0.164 | -4.067 | -1.240 | 0.010 |
| | 1.3°, 90° | -0.865 | 0.159 | -5.433 | -1.611 | < .001 |
| | 4.8°, 90° | -0.428 | 0.164 | -2.617 | -0.798 | 1.000 |
| 4.8°, 0° | .096°, 30° | -0.016 | 0.164 | -0.100 | -0.030 | 1.000 |
| | .4°, 30° | -0.005 | 0.164 | -0.028 | -0.008 | 1.000 |
| | 1.3°, 30° | 0.230 | 0.164 | 1.406 | 0.429 | 1.000 |
| | 4.8°, 30° | 0.065 | 0.159 | 0.411 | 0.122 | 1.000 |
| | .096°, 60° | -0.155 | 0.164 | -0.950 | -0.290 | 1.000 |
| | .4°, 60° | -0.155 | 0.164 | -0.950 | -0.290 | 1.000 |
| | 1.3°, 60° | -0.115 | 0.164 | -0.700 | -0.213 | 1.000 |
| | 4.8°, 60° | 0.042 | 0.159 | 0.263 | 0.078 | 1.000 |
| | .096°, 90° | -0.087 | 0.164 | -0.533 | -0.163 | 1.000 |
| | .4°, 90° | -0.542 | 0.164 | -3.312 | -1.010 | 0.145 |
| | 1.3°, 90° | -0.741 | 0.164 | -4.528 | -1.381 | 0.002 |
| | 4.8°, 90° | -0.305 | 0.159 | -1.914 | -0.568 | 1.000 |
| .096°, 30° | .4°, 30° | 0.012 | 0.152 | 0.078 | 0.022 | 1.000 |
| | 1.3°, 30° | 0.246 | 0.152 | 1.618 | 0.459 | 1.000 |
| | 4.8°, 30° | 0.082 | 0.152 | 0.537 | 0.152 | 1.000 |
| | .096°, 60° | -0.139 | 0.159 | -0.874 | -0.259 | 1.000 |
| | .4°, 60° | -0.139 | 0.164 | -0.850 | -0.259 | 1.000 |
| | 1.3°, 60° | -0.098 | 0.164 | -0.600 | -0.183 | 1.000 |

| | | | | | | |
|---|---|---|---|---|---|---|
| | 4.8°, 60° | 0.058 | 0.164 | 0.356 | 0.108 | 1.000 |
| | .096°, 90° | -0.071 | 0.159 | -0.446 | -0.132 | 1.000 |
| | .4°, 90° | -0.525 | 0.164 | -3.212 | -0.979 | 0.200 |
| | 1.3°, 90° | -0.725 | 0.164 | -4.428 | -1.350 | 0.002 |
| | 4.8°, 90° | -0.288 | 0.164 | -1.761 | -0.537 | 1.000 |
| .4°, 30° | 1.3°, 30° | 0.235 | 0.152 | 1.541 | 0.437 | 1.000 |
| | 4.8°, 30° | 0.070 | 0.152 | 0.460 | 0.130 | 1.000 |
| | .096°, 60° | -0.151 | 0.164 | -0.922 | -0.281 | 1.000 |
| | .4°, 60° | -0.151 | 0.159 | -0.948 | -0.281 | 1.000 |
| | 1.3°, 60° | -0.110 | 0.164 | -0.672 | -0.205 | 1.000 |
| | 4.8°, 60° | 0.046 | 0.164 | 0.283 | 0.086 | 1.000 |
| | .096°, 90° | -0.083 | 0.164 | -0.506 | -0.154 | 1.000 |
| | .4°, 90° | -0.537 | 0.159 | -3.376 | -1.001 | 0.121 |
| | 1.3°, 90° | -0.736 | 0.164 | -4.501 | -1.372 | 0.002 |
| | 4.8°, 90° | -0.300 | 0.164 | -1.834 | -0.559 | 1.000 |
| 1.3°, 30° | 4.8°, 30° | -0.165 | 0.152 | -1.081 | -0.307 | 1.000 |
| | .096°, 60° | -0.385 | 0.164 | -2.356 | -0.718 | 1.000 |
| | .4°, 60° | -0.385 | 0.164 | -2.356 | -0.718 | 1.000 |
| | 1.3°, 60° | -0.345 | 0.159 | -2.165 | -0.642 | 1.000 |
| | 4.8°, 60° | -0.188 | 0.164 | -1.150 | -0.351 | 1.000 |
| | .096°, 90° | -0.317 | 0.164 | -1.939 | -0.591 | 1.000 |
| | .4°, 90° | -0.772 | 0.164 | -4.717 | -1.438 | < .001 |
| | 1.3°, 90° | -0.971 | 0.159 | -6.101 | -1.809 | < .001 |
| | 4.8°, 90° | -0.535 | 0.164 | -3.267 | -0.996 | 0.167 |
| 4.8°, 30° | .096°, 60° | -0.221 | 0.164 | -1.350 | -0.412 | 1.000 |
| | .4°, 60° | -0.221 | 0.164 | -1.350 | -0.412 | 1.000 |
| | 1.3°, 60° | -0.180 | 0.164 | -1.100 | -0.335 | 1.000 |
| | 4.8°, 60° | -0.024 | 0.159 | -0.149 | -0.044 | 1.000 |
| | .096°, 90° | -0.153 | 0.164 | -0.933 | -0.285 | 1.000 |
| | .4°, 90° | -0.607 | 0.164 | -3.712 | -1.132 | 0.037 |
| | 1.3°, 90° | -0.806 | 0.164 | -4.929 | -1.503 | < .001 |
| | 4.8°, 90° | -0.370 | 0.159 | -2.325 | -0.690 | 1.000 |
| .096°, 60° | .4°, 60° | < -0.001 | 0.152 | < -0.001 | < -0.001 | 1.000 |
| | 1.3°, 60° | 0.041 | 0.152 | 0.269 | 0.076 | 1.000 |
| | 4.8°, 60° | 0.197 | 0.152 | 1.296 | 0.368 | 1.000 |
| | .096°, 90° | 0.068 | 0.159 | 0.428 | 0.127 | 1.000 |
| | .4°, 90° | -0.386 | 0.164 | -2.361 | -0.720 | 1.000 |
| | 1.3°, 90° | -0.585 | 0.164 | -3.578 | -1.091 | 0.059 |
| | 4.8°, 90° | -0.149 | 0.164 | -0.911 | -0.278 | 1.000 |
| .4°, 60° | 1.3°, 60° | 0.041 | 0.152 | 0.269 | 0.076 | 1.000 |
| | 4.8°, 60° | 0.197 | 0.152 | 1.296 | 0.368 | 1.000 |
| | .096°, 90° | 0.068 | 0.164 | 0.417 | 0.127 | 1.000 |
| | .4°, 90° | -0.386 | 0.159 | -2.428 | -0.720 | 1.000 |
| | 1.3°, 90° | -0.585 | 0.164 | -3.578 | -1.091 | 0.059 |
| | 4.8°, 90° | -0.149 | 0.164 | -0.911 | -0.278 | 1.000 |
| 1.3°, 60° | 4.8°, 60° | 0.156 | 0.152 | 1.027 | 0.291 | 1.000 |

| | | | | | | |
|---|---|---|---|---|---|---|
| | .096°, 90° | 0.027 | 0.164 | 0.167 | 0.051 | 1.000 |
| | .4°, 90° | -0.427 | 0.164 | -2.612 | -0.796 | 1.000 |
| | 1.3°, 90° | -0.626 | 0.159 | -3.936 | -1.167 | 0.017 |
| | 4.8°, 90° | -0.190 | 0.164 | -1.161 | -0.354 | 1.000 |
| 4.8°, 60° | .096°, 90° | -0.129 | 0.164 | -0.789 | -0.241 | 1.000 |
| | .4°, 90° | -0.584 | 0.164 | -3.567 | -1.088 | 0.061 |
| | 1.3°, 90° | -0.783 | 0.164 | -4.784 | -1.459 | < .001 |
| | 4.8°, 90° | -0.346 | 0.159 | -2.176 | -0.645 | 1.000 |
| .096°, 90° | .4°, 90° | -0.455 | 0.152 | -2.986 | -0.847 | 0.412 |
| | 1.3°, 90° | -0.654 | 0.152 | -4.294 | -1.218 | 0.004 |
| | 4.8°, 90° | -0.217 | 0.152 | -1.427 | -0.405 | 1.000 |
| .4°, 90° | 1.3°, 90° | -0.199 | 0.152 | -1.308 | -0.371 | 1.000 |
| | 4.8°, 90° | 0.237 | 0.152 | 1.559 | 0.442 | 1.000 |
| 1.3°, 90° | 4.8°, 90° | 0.436 | 0.152 | 2.866 | 0.813 | 0.590 |

*Note.* P-value adjusted for comparing a family of 120 estimates.

Experiment 4

**Supplementary Table 7**

*Post hoc tests of hue angle for $I_1$ (14.66 Hz) in Experiment 4*

| | | Mean Difference | SE | t | Cohen's d | $p_{bonf}$ |
|---|---|---|---|---|---|---|
| 0° | 30° | 0.255 | 0.114 | 2.223 | 0.664 | 1.000 |
| | 60° | 0.671 | 0.114 | 5.866 | 1.751 | < .001 |
| | 90° | 1.021 | 0.114 | 8.920 | 2.662 | < .001 |
| | 120° | 0.756 | 0.114 | 6.604 | 1.971 | < .001 |
| | 150° | 0.552 | 0.114 | 4.822 | 1.439 | < .001 |
| | 180° | 0.450 | 0.114 | 3.935 | 1.174 | 0.010 |
| | 210° | 0.677 | 0.114 | 5.915 | 1.765 | < .001 |
| | 240° | 0.928 | 0.114 | 8.109 | 2.420 | < .001 |
| | 270° | 1.055 | 0.114 | 9.215 | 2.750 | < .001 |
| | 300° | 0.489 | 0.114 | 4.268 | 1.274 | 0.003 |
| | 330° | 0.087 | 0.114 | 0.756 | 0.226 | 1.000 |
| 30° | 60° | 0.417 | 0.114 | 3.642 | 1.087 | 0.029 |
| | 90° | 0.767 | 0.114 | 6.697 | 1.999 | < .001 |
| | 120° | 0.502 | 0.114 | 4.381 | 1.308 | 0.002 |
| | 150° | 0.297 | 0.114 | 2.598 | 0.776 | 0.712 |
| | 180° | 0.196 | 0.114 | 1.711 | 0.511 | 1.000 |
| | 210° | 0.423 | 0.114 | 3.692 | 1.102 | 0.024 |
| | 240° | 0.674 | 0.114 | 5.886 | 1.757 | < .001 |
| | 270° | 0.800 | 0.114 | 6.992 | 2.087 | < .001 |
| | 300° | 0.234 | 0.114 | 2.045 | 0.610 | 1.000 |
| | 330° | -0.168 | 0.114 | -1.467 | -0.438 | 1.000 |
| 60° | 90° | 0.350 | 0.114 | 3.054 | 0.912 | 0.191 |
| | 120° | 0.085 | 0.114 | 0.739 | 0.221 | 1.000 |
| | 150° | -0.120 | 0.114 | -1.044 | -0.312 | 1.000 |
| | 180° | -0.221 | 0.114 | -1.931 | -0.576 | 1.000 |
| | 210° | 0.006 | 0.114 | 0.050 | 0.015 | 1.000 |

| | | | | | | |
|---|---|---|---|---|---|---|
| | 240° | 0.257 | 0.114 | 2.244 | 0.670 | 1.000 |
| | 270° | 0.383 | 0.114 | 3.349 | 1.000 | 0.076 |
| | 300° | -0.183 | 0.114 | -1.598 | -0.477 | 1.000 |
| | 330° | -0.585 | 0.114 | -5.110 | -1.525 | < .001 |
| 90° | 120° | -0.265 | 0.114 | -2.316 | -0.691 | 1.000 |
| | 150° | -0.469 | 0.114 | -4.099 | -1.223 | 0.006 |
| | 180° | -0.571 | 0.114 | -4.985 | -1.488 | < .001 |
| | 210° | -0.344 | 0.114 | -3.005 | -0.897 | 0.222 |
| | 240° | -0.093 | 0.114 | -0.811 | -0.242 | 1.000 |
| | 270° | 0.034 | 0.114 | 0.295 | 0.088 | 1.000 |
| | 300° | -0.533 | 0.114 | -4.652 | -1.389 | < .001 |
| | 330° | -0.935 | 0.114 | -8.164 | -2.437 | < .001 |
| 120° | 150° | -0.204 | 0.114 | -1.783 | -0.532 | 1.000 |
| | 180° | -0.306 | 0.114 | -2.670 | -0.797 | 0.585 |
| | 210° | -0.079 | 0.114 | -0.689 | -0.206 | 1.000 |
| | 240° | 0.172 | 0.114 | 1.505 | 0.449 | 1.000 |
| | 270° | 0.299 | 0.114 | 2.611 | 0.779 | 0.689 |
| | 300° | -0.268 | 0.114 | -2.337 | -0.697 | 1.000 |
| | 330° | -0.670 | 0.114 | -5.849 | -1.746 | < .001 |
| 150° | 180° | -0.102 | 0.114 | -0.887 | -0.265 | 1.000 |
| | 210° | 0.125 | 0.114 | 1.094 | 0.326 | 1.000 |
| | 240° | 0.376 | 0.114 | 3.288 | 0.981 | 0.092 |
| | 270° | 0.503 | 0.114 | 4.394 | 1.311 | 0.002 |
| | 300° | -0.063 | 0.114 | -0.554 | -0.165 | 1.000 |
| | 330° | -0.465 | 0.114 | -4.066 | -1.213 | 0.006 |
| 180° | 210° | 0.227 | 0.114 | 1.981 | 0.591 | 1.000 |
| | 240° | 0.478 | 0.114 | 4.175 | 1.246 | 0.004 |
| | 270° | 0.605 | 0.114 | 5.280 | 1.576 | < .001 |
| | 300° | 0.038 | 0.114 | 0.333 | 0.099 | 1.000 |
| | 330° | -0.364 | 0.114 | -3.179 | -0.949 | 0.130 |
| 210° | 240° | 0.251 | 0.114 | 2.194 | 0.655 | 1.000 |
| | 270° | 0.378 | 0.114 | 3.300 | 0.985 | 0.089 |
| | 300° | -0.189 | 0.114 | -1.647 | -0.492 | 1.000 |
| | 330° | -0.591 | 0.114 | -5.159 | -1.540 | < .001 |
| 240° | 270° | 0.127 | 0.114 | 1.106 | 0.330 | 1.000 |
| | 300° | -0.440 | 0.114 | -3.841 | -1.147 | 0.014 |
| | 330° | -0.842 | 0.114 | -7.353 | -2.195 | < .001 |
| 270° | 300° | -0.566 | 0.114 | -4.947 | -1.477 | < .001 |
| | 330° | -0.968 | 0.114 | -8.459 | -2.525 | < .001 |
| 300° | 330° | -0.402 | 0.114 | -3.512 | -1.048 | 0.044 |

*Note.* P-value adjusted for comparing a family of 66 estimates.

**Supplementary Table 8**

*Post hoc tests of hue angle for $F_B$ (8 Hz) in Experiment 4*

| | | Mean Difference | SE | t | Cohen's d | $p_{bonf}$ |
|---|---|---|---|---|---|---|
| 0° | 30° | 0.074 | 0.125 | 0.590 | 0.168 | 1.000 |

| | | | | | | |
|---|---|---|---|---|---|---|
| | 60° | -0.303 | 0.125 | -2.416 | -0.689 | 1.000 |
| | 90° | -0.419 | 0.125 | -3.338 | -0.953 | 0.078 |
| | 120° | -0.570 | 0.125 | -4.544 | -1.297 | 0.001 |
| | 150° | -0.587 | 0.125 | -4.685 | -1.337 | < .001 |
| | 180° | -0.733 | 0.125 | -5.849 | -1.669 | < .001 |
| | 210° | -0.721 | 0.125 | -5.751 | -1.641 | < .001 |
| | 240° | -0.541 | 0.125 | -4.311 | -1.230 | 0.003 |
| | 270° | -0.392 | 0.125 | -3.125 | -0.892 | 0.154 |
| | 300° | -0.132 | 0.125 | -1.055 | -0.301 | 1.000 |
| | 330° | -0.083 | 0.125 | -0.661 | -0.189 | 1.000 |
| 30° | 60° | -0.377 | 0.125 | -3.006 | -0.858 | 0.222 |
| | 90° | -0.493 | 0.125 | -3.928 | -1.121 | 0.010 |
| | 120° | -0.644 | 0.125 | -5.134 | -1.465 | < .001 |
| | 150° | -0.661 | 0.125 | -5.275 | -1.505 | < .001 |
| | 180° | -0.807 | 0.125 | -6.439 | -1.838 | < .001 |
| | 210° | -0.795 | 0.125 | -6.341 | -1.810 | < .001 |
| | 240° | -0.615 | 0.125 | -4.901 | -1.399 | < .001 |
| | 270° | -0.466 | 0.125 | -3.715 | -1.060 | 0.022 |
| | 300° | -0.206 | 0.125 | -1.645 | -0.469 | 1.000 |
| | 330° | -0.157 | 0.125 | -1.251 | -0.357 | 1.000 |
| 60° | 90° | -0.116 | 0.125 | -0.923 | -0.263 | 1.000 |
| | 120° | -0.267 | 0.125 | -2.128 | -0.607 | 1.000 |
| | 150° | -0.285 | 0.125 | -2.269 | -0.648 | 1.000 |
| | 180° | -0.430 | 0.125 | -3.433 | -0.980 | 0.058 |
| | 210° | -0.418 | 0.125 | -3.336 | -0.952 | 0.079 |
| | 240° | -0.238 | 0.125 | -1.896 | -0.541 | 1.000 |
| | 270° | -0.089 | 0.125 | -0.709 | -0.202 | 1.000 |
| | 300° | 0.171 | 0.125 | 1.361 | 0.388 | 1.000 |
| | 330° | 0.220 | 0.125 | 1.755 | 0.501 | 1.000 |
| 90° | 120° | -0.151 | 0.125 | -1.205 | -0.344 | 1.000 |
| | 150° | -0.169 | 0.125 | -1.347 | -0.384 | 1.000 |
| | 180° | -0.315 | 0.125 | -2.511 | -0.717 | 0.902 |
| | 210° | -0.303 | 0.125 | -2.413 | -0.689 | 1.000 |
| | 240° | -0.122 | 0.125 | -0.973 | -0.278 | 1.000 |
| | 270° | 0.027 | 0.125 | 0.213 | 0.061 | 1.000 |
| | 300° | 0.286 | 0.125 | 2.283 | 0.652 | 1.000 |
| | 330° | 0.336 | 0.125 | 2.677 | 0.764 | 0.573 |
| 120° | 150° | -0.018 | 0.125 | -0.141 | -0.040 | 1.000 |
| | 180° | -0.164 | 0.125 | -1.305 | -0.373 | 1.000 |
| | 210° | -0.151 | 0.125 | -1.208 | -0.345 | 1.000 |
| | 240° | 0.029 | 0.125 | 0.232 | 0.066 | 1.000 |
| | 270° | 0.178 | 0.125 | 1.419 | 0.405 | 1.000 |
| | 300° | 0.437 | 0.125 | 3.489 | 0.996 | 0.048 |
| | 330° | 0.487 | 0.125 | 3.883 | 1.108 | 0.012 |
| 150° | 180° | -0.146 | 0.125 | -1.164 | -0.332 | 1.000 |
| | 210° | -0.134 | 0.125 | -1.066 | -0.304 | 1.000 |

| | | | | | | |
|---|---|---|---|---|---|---|
| | 240° | 0.047 | 0.125 | 0.374 | 0.107 | 1.000 |
| | 270° | 0.196 | 0.125 | 1.560 | 0.445 | 1.000 |
| | 300° | 0.455 | 0.125 | 3.630 | 1.036 | 0.030 |
| | 330° | 0.505 | 0.125 | 4.024 | 1.148 | 0.007 |
| 180° | 210° | 0.012 | 0.125 | 0.098 | 0.028 | 1.000 |
| | 240° | 0.193 | 0.125 | 1.538 | 0.439 | 1.000 |
| | 270° | 0.342 | 0.125 | 2.724 | 0.777 | 0.503 |
| | 300° | 0.601 | 0.125 | 4.794 | 1.368 | < .001 |
| | 330° | 0.650 | 0.125 | 5.188 | 1.481 | < .001 |
| 210° | 240° | 0.181 | 0.125 | 1.440 | 0.411 | 1.000 |
| | 270° | 0.329 | 0.125 | 2.626 | 0.750 | 0.660 |
| | 300° | 0.589 | 0.125 | 4.696 | 1.340 | < .001 |
| | 330° | 0.638 | 0.125 | 5.090 | 1.453 | < .001 |
| 240° | 270° | 0.149 | 0.125 | 1.186 | 0.339 | 1.000 |
| | 300° | 0.408 | 0.125 | 3.256 | 0.929 | 0.102 |
| | 330° | 0.458 | 0.125 | 3.650 | 1.042 | 0.028 |
| 270° | 300° | 0.260 | 0.125 | 2.070 | 0.591 | 1.000 |
| | 330° | 0.309 | 0.125 | 2.464 | 0.703 | 1.000 |
| 300° | 330° | 0.049 | 0.125 | 0.394 | 0.112 | 1.000 |

*Note.* P-value adjusted for comparing a family of 66 estimates.

**2. Treatment of eye-based artefacts**

We have used two alternative approaches to treatment of eye-based artefacts (blinks and eye movements) in our analysis pipeline. In Experiments 1 and 4, no inspection of trials for eye-based artefacts was performed prior to further analysis. In Experiments 2 and 3, all trials were visually inspected for artefacts and were removed from further analysis if eye-based artefacts were present. If the artefacts occurred on less than 20% of total trials in the condition, complete trials were removed. In cases where the artefacts were present in 20% or more of the trials, independent component analysis (ICA) was performed and the identified components corresponding to eye-based artefacts were removed.

We present both approaches as we found no meaningful difference in final SNR between the two approaches. We inspected the effect of alternative approaches on final SNR on data from the S/(L+M) condition in Experiment 1. The data was analyzed and SNR at $I_1$ (14.66 Hz) and $F_B$ (8 Hz) components extracted for each of the two approaches.

As visualized in Supplementary Figure 1, the difference in SNR between the two approaches was minimal.

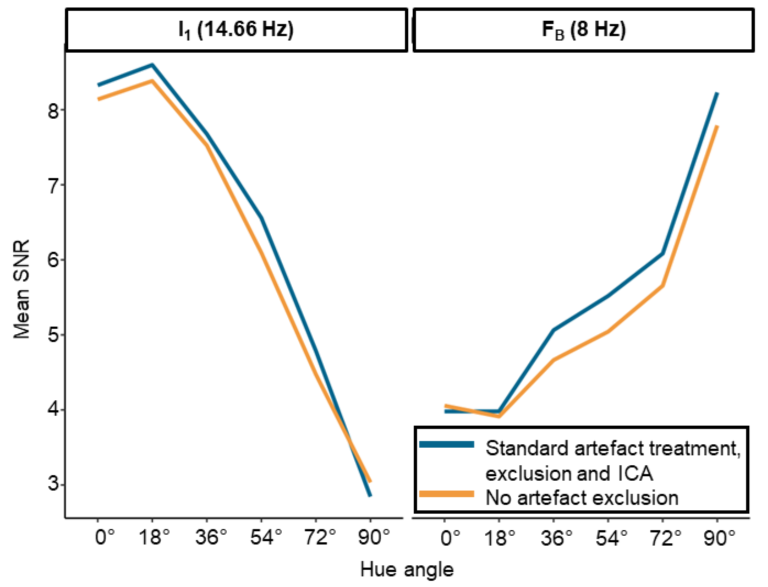


**Supplementary Figure 1. Signal-to-noise ratios for the two ocular artefact treatments. The data in the example are from Experiment 1, for the S/(L+M) axis condition which was analyzed with both pipelines – either with ocular artefacts manually checked and excluded (blue line), or with no exclusions made (orange line). The outcomes are presented as a function of hue angle difference between fixed and variable chromaticity and separately for the interaction component (left, $I_1$) and base frequency (right, $F_B$).**

In Supplementary Figure 2 we visualize the difference between the two approaches. This was derived by expressing the SNR for the pipeline without blink exclusions as percentage of the SNR for the pipeline with blink exclusions (as applied in Experiment 2 and 3). The difference score presented in Supplementary Figure 3 is expressed as a percentage. If the two approaches completely matched SNR, the difference between them should be 0. Any positive difference shows higher SNR in for the pipeline where no artefacts were excluded, and a negative difference a higher SNR in standard blink exclusion pipeline. For the S/(L+M) condition in Experiment 1, the absolute difference has not surpassed 10% in any condition. We have thus deemed the two approaches to eye-based artifact removal as producing similar outcomes.

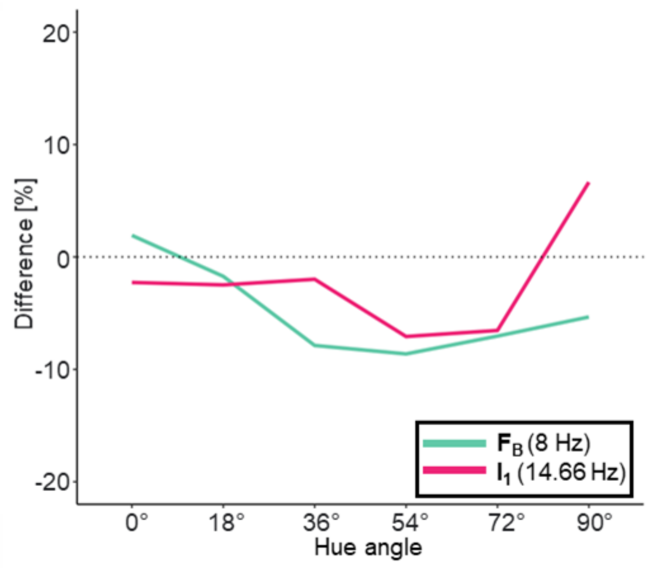


**Supplementary Figure 2. Difference in SNR between ocular artefact treatment in data processing. The difference in SNR between the no exclusions pipeline expressed as percentage of the SNR in the standard exclusion pipeline. The data are from Experiment 1, S/(L+M) axis condition. The outcomes are presented as a function of hue angle difference between fixed and variable chromaticity and separately for the intermodulation component (pink, $I_1$) and base frequency (aqua, $F_B$). The dotted line denotes the point of no difference between pipelines (0% difference).**

**3. Scalp maps for Experiments 1 and 4**

We extracted signals at all scalp locations for Experiments 1 and 4. We present these as scalp heatmap plots in Supplementary Figures 3 and 4, respectively.

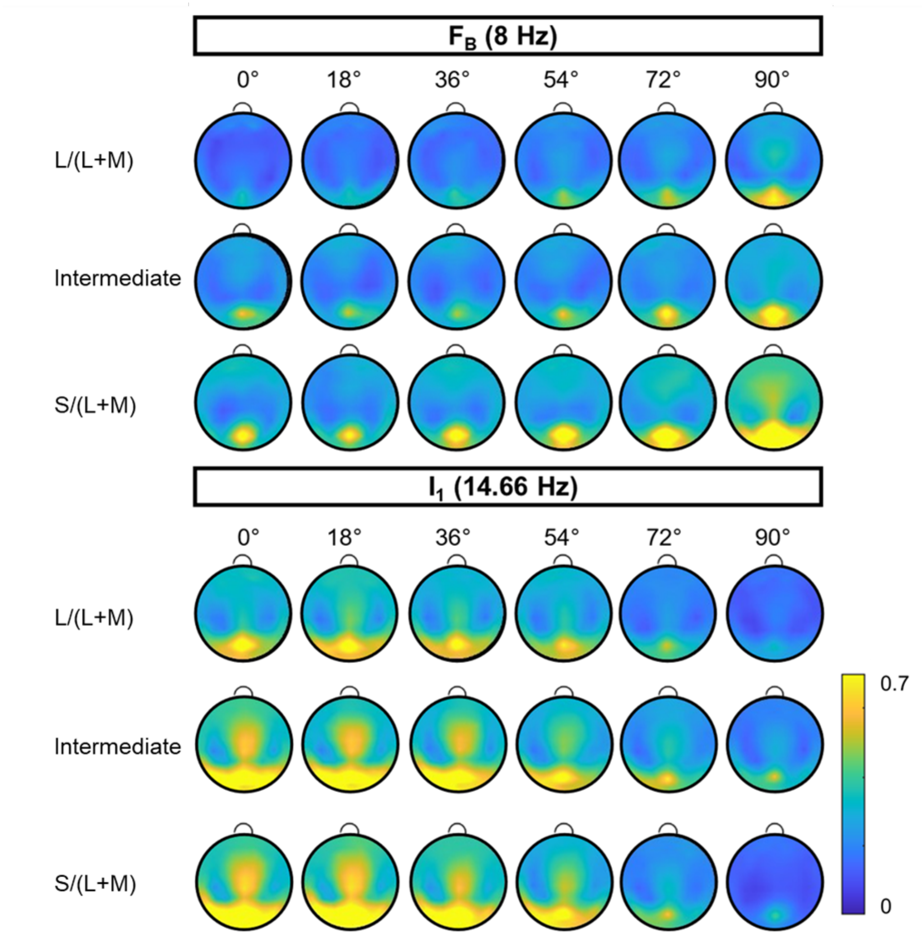


**Supplementary Figure 3. Heatmaps of average amplitudes within Experiment 1. The amplitudes reflect the average across observers.**

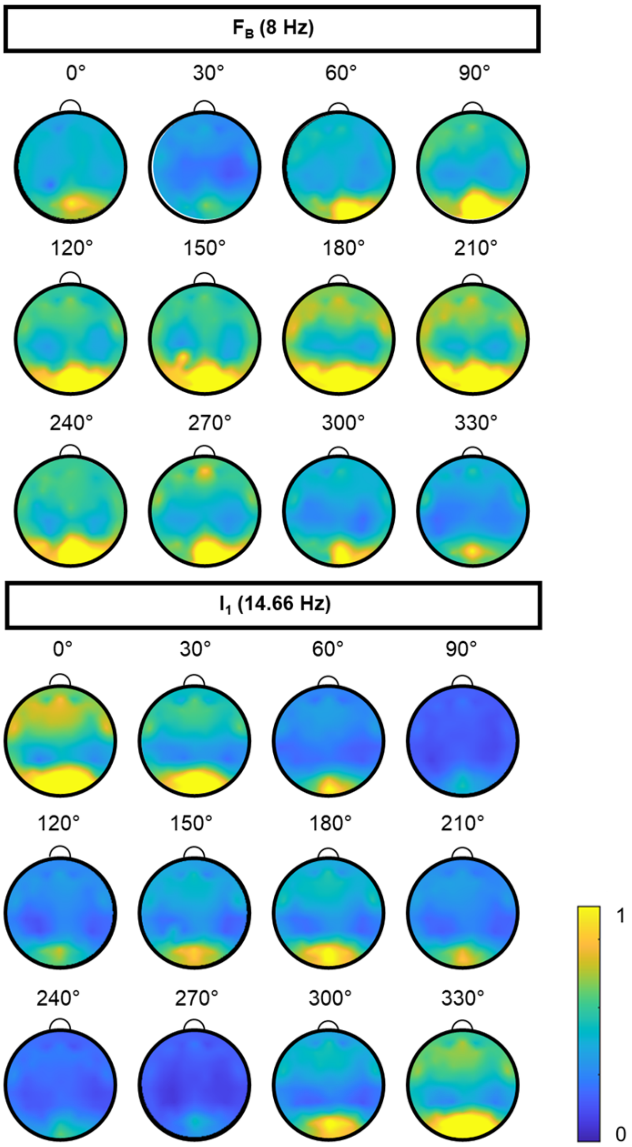


**Supplementary Figure 4. Heatmaps of average amplitudes within Experiment 4. The amplitudes reflect the average across observers.**

**4. Tuning across intermodulation components**

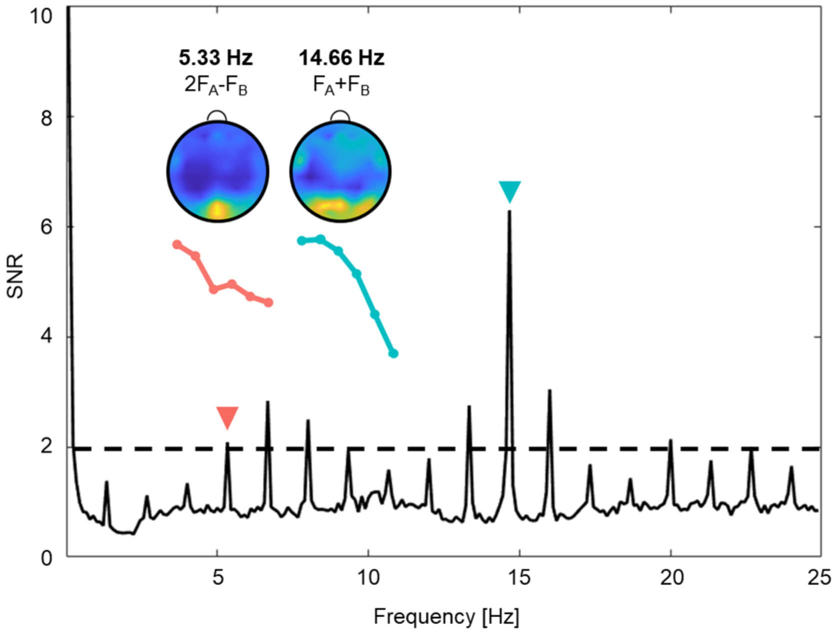


**Supplementary Figure 5. Signal heatmaps and tuning functions at intermodulation components surpassing 2 SNR in Experiment 1. SNR is presented as a function of Frequency in Hz with a range of 0 – 25. The two components and their source (nature of intermodulation between $F_A$ and $F_B$) are labelled above the individual heatmap and tuning function plots. The scales of individual heatmaps and tuning function plots are scaled to match in area under curve.**

**5. Differences in tuning functions between cortical locations**

One of the limitations of EEG as a neuroimaging method is its coarse spatial resolution. As an additional analysis, we wanted to check if cortical color functions change between cortical locations. We chose to separate the most posterior part of the scalp into three subgroups, each encompassing one set of electrodes, from the most posterior occipital cluster, through a parietal-occipital cluster to a parietal cluster, most anterior of the three clusters (Supplementary Figure 5c).

For Experiments 1 and 4, data from all electrodes were extracted at $I_1$ (14.66 Hz) and $F_B$ (8 Hz). The electrode signals were inspected in the same way as for the main analyses, described in the general methods section. A single electrode exclusion (PO3) was made for one observer in Experiment 4.

The amplitude was averaged across observers and electrodes in the sub-cluster (O, PO and P for occipital, parietal-occipital and parietal respectively). In order to compare the shapes of the tuning functions between clusters, the amplitudes were scaled to match in area under the curve. The correspondence in function shape is demonstrated in Supplementary Figure 6 panels a (Experiment 1) and b (Experiment 4). We interpret the matching shape as a transitive feature of EEG signals where strong signal emerging from one location can also be recorded in proximal channels. Consequently, we find no difference in tuning function shapes between more posterior and more anterior sites and conclude that imaging methods with more precise spatial resolution (such as fMRI) would be neded to determine the difference in tuning between different cortical sites.

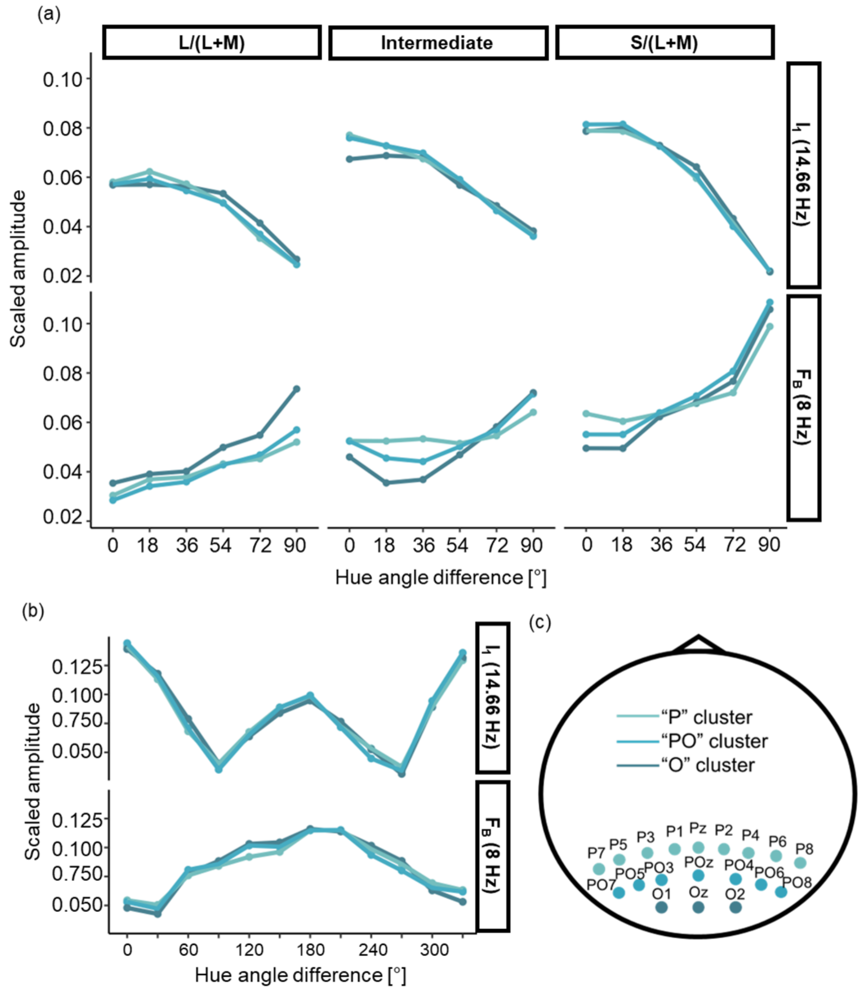


**Supplementary Figure 6. Overview of tuning functions at different locations. Panel (a) shows results from Experiment 1 and panel (b) shows results from Experiment 4. Each tuning function is scaled to match the area under the curve. A schematic representation of approximate scalp locations is presented in panel (c), together with a color legend for the figure.**

### 6. Stimulus signal check

During stimulus presentation, a photodiode was placed near the top of the display. The sensor was located to capture the edge between two checks in the checkerboard.

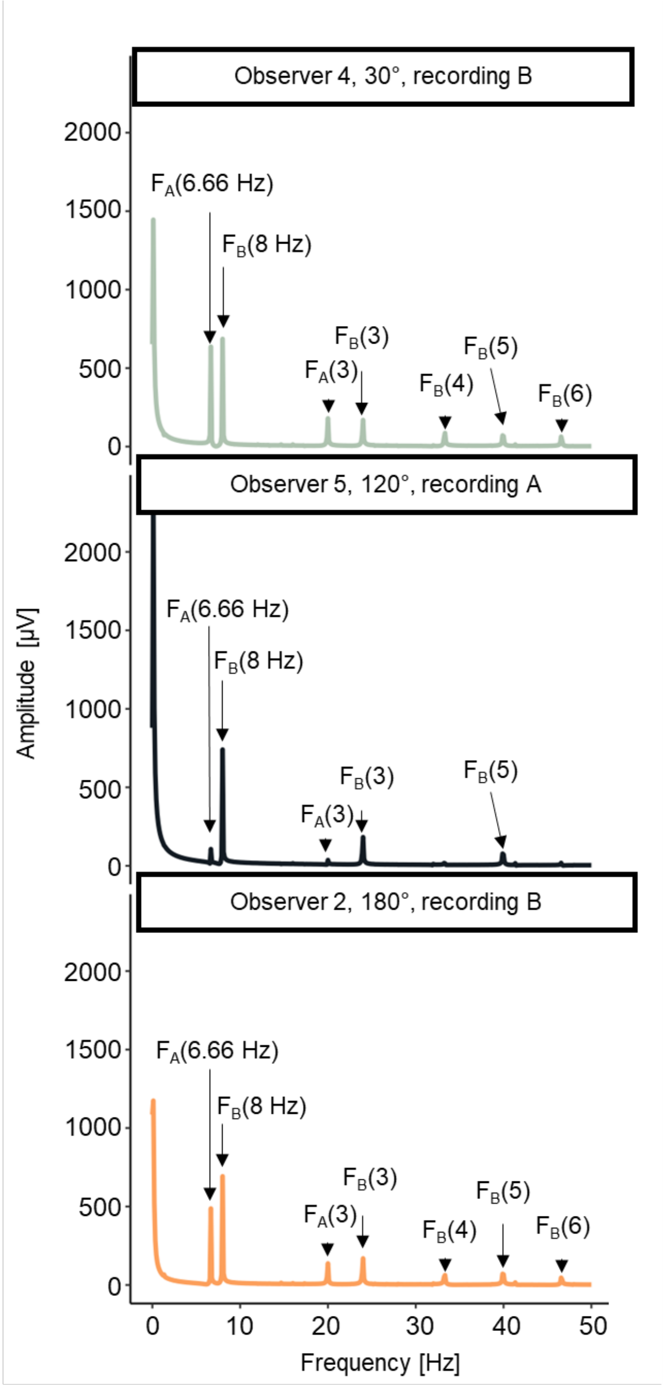


**Supplementary Figure 7. Frequency profile from three randomly selected example photodiode recordings in Experiment 4. Each presents the photodiode profile for a specific recording for a specific observer, for a specific condition, summarized above each plot. The visible peaks in up to 50 Hz are labelled within each plot.**
